# Mechanoimmunological Control of Metastatic Site Selection

**DOI:** 10.1101/2025.05.10.653256

**Authors:** Yassmin A. Elbanna, Maria Tello-Lafoz, Aliya Holland, Ye Zhang, Jun-Goo Kwak, Zhenghan Wang, Alexandrina Yakimov, Myra Dada, Samuel Vayner, Sarah M. Duquette, Young Hun Kim, Tejus A. Bale, Benjamin Y. Winer, Kenny K. H. Yu, Joan Massagué, Jungwoo Lee, Ori Barzilai, Scott R. Manalis, Morgan Huse

## Abstract

Cancer cells alter their mechanical properties in response to the rigidity of their environment. Here, we explored the implications of this environmental mechanosensing for anti-tumor immunosurveillance using single cell biophysical profiling and metastasis models. Cancer cells stiffened in more rigid environments, a biophysical change that sensitized them to cytotoxic lymphocytes. In immunodeficient mice, this behavior manifested in the outgrowth of stiffer metastatic cells in the rigid bone than in the soft lung, while in immunocompetent hosts, it led to preferential elimination of stiffer cancer cells and suppression of bone metastasis. Environmentally-induced cell stiffening and immune sensitization both required Osteopontin, a secreted glycoprotein that is upregulated during bone colonization. Analysis of patient metastases spanning mechanically distinct tissues revealed associations between environmental rigidity, immune infiltration, and cancer cell stiffness consistent with mechanically driven immunosurveillance. These results demonstrate how environmental mechanosensing modulates anti-tumor immunity and suggest a mechanoimmunological basis for metastatic site selection.

## Introduction

Metastasis accounts for 90% of cancer-related deaths (*1*), underscoring the urgent need to understand the mechanisms underlying metastatic progression and colonization. A defining trait of metastasis is its positional diversity, with each type of disease colonizing an array of target organs that vary widely in their molecular composition and tissue architecture (*2-4*). Much is now known about the cellular and biochemical features of distinct metastatic niches and how they influence cancer cell colonization. By contrast, the interplay between the mechanical properties of these microenvironments and the cancer cells within them remains poorly understood. Rigidity, defined as the resistance of material to deformation, is particularly variable between metastatic niches, with organs like the lung and brain being very soft and mineralized bone approaching the rigidity of steel (*5*).

Adherent cell types, including many cancer cells, engage in continuous biomechanical crosstalk with their tissue environment (*6-9*). This process, termed mechanoreciprocity, is initiated by cytoskeletally-derived force exertion against the extracellular matrix (ECM) and other cells in the surrounding milieu. Physical probing of this kind facilitates the activation of mechanosensitive receptors, such as integrins, which in turn transduce signals that stimulate cancer cell proliferation and influence cell morphology, polarity, and gene expression. Concomitantly, cancer cells alter the mechanics of their environment by both physically deforming it and by releasing additional ECM components. An important consequence of mechanoreciprocity is that the responding cell mimics the physical properties of its surroundings, becoming stiffer in more rigid environments and softer in more pliable ones (*10-13*). One would therefore expect that metastatic tumor cells (MTCs) occupying rigid niches like the bone would be stiffer than those in more pliable tissues. The importance of this type of mechanical equilibration for the efficacy of metastatic colonization is not known.

Metastatic progression is antagonized by cytotoxic lymphocytes, comprising CD8^+^ cytotoxic T lymphocytes (CTLs) and natural killer (NK) cells (*14, 15*). CTLs attack transformed target cells expressing neoantigens (*16*), while NK cells attack targets that express stress-induced ligands or exhibit indices of immune recognition (e.g. antibody opsonization) (*17*). Target cell engagement initiates the formation of a stereotyped cell-cell interaction, known as the immune synapse, into which the cytotoxic lymphocyte secretes a toxic mixture of granzyme proteases and the pore forming protein perforin to elicit programmed cell death (*18*). The importance of this pathway for anti-tumor immunity is highlighted by the clinical success of treatment modalities, such as immune checkpoint blockade (ICB), that function by boosting the activity of cytotoxic lymphocytes(*19*).

In recent years, it has become clear that immune synapses respond not only to the biochemical composition of the target cell surface but also to its mechanical properties (*20*). The basis for this mechanosensitive behavior is thought to arise from the activating immunoreceptors responsible for synapse formation. A number of these proteins, including the T cell antigen receptor (TCR), certain activating NK receptors, and integrins like LFA-1 (for Lymphocyte Function Associated antigen-1, α_L_β_2_), only achieve optimal ligand binding and signaling when placed under tension (*21-23*). This property imposes physical demands on the surface of the target cell, which must be rigid enough to counterbalance the load placed on these receptors and their associated ligands. Hence, stiffer surfaces bearing stimulatory ligands induce stronger lymphocyte activation than softer surfaces coated with the same proteins (*24-27*). We and others have found that this property leads to enhanced destruction of stiffer cancer cells by cytotoxic lymphocytes (*28-30*). This biophysically-directed killing response, called mechanosurveillance, appears to be particularly important during metastasis, when cancer cells lack the protection of a highly immunosuppressive microenvironment (*28*).

In the present study, we explored the implications of mechanosurveillance and mechanoreciprocity for metastatic progression in biophysically disparate organs. We hypothesized that cancer cells colonizing rigid *in vivo* environments would stiffen mechanoreciprocally and that this would sensitize them to destruction by mechanosensitive cytotoxic lymphocytes. Consistent with this prediction, we found that cytotoxic lymphocytes strongly suppressed bone colonization in murine models of metastasis while also selectively eliminating mechanically stiffer cancer cells. We further identified Opn, a secreted ECM protein, as a critical mediator of environmentally-induced cancer cell stiffening and immune targeting. Finally, we validated the relevance of environmental rigidity and immune stiffness sensing for human metastasis using paired patient samples from mechanically distinct locations. Collectively, these results demonstrate how the interplay between mechanoreciprocity and mechanosurveillance can effectively dictate not only the amount of metastatic burden but also the sites where outgrowth occurs.

## Results

### Environmental rigidity controls the stiffness of cancer cells and their sensitivity to cytotoxic lymphocytes

To explore the effects of environmental rigidity on the stiffness and immune vulnerability of highly metastatic cells, we cultured B16F10 melanoma cells on fibronectin-coated substrates of varying rigidity, ranging from soft polyacrylamide hydrogels (8 kPa Young’s Modulus) to stiffer hydrogels (50 kPa) and tissue culture plastic (∼1 GPa). After 24 hours, cells were replated on glass and their stiffness measured by atomic force microscopy (AFM) (Fig. 1A). We observed a progressive increase in cell stiffness that reflected the rigidity of the initial substrate; approximately 40% of cells cultured on plastic exceeded 3 kPa in stiffness, whereas only ∼20% and ∼10% of cells cultured on 50 kPa and 8 kPa hydrogel, respectively, reached this threshold (Fig. 1B). These differences fell short of statistical significance, however, likely due to insufficient sample size. To address this issue, we used an alternative approach based on the suspended microchannel resonator (SMR), a microfluidic device in which suspension cells flow through a hairpin-shaped microchannel embedded a vibrating cantilever (Fig. 1C). The vibration creates a standing acoustic wave within the channel, and as cells flow through the node of the cantilever, their interaction with the wave induces shifts in its vibrational frequency. These shifts are used to calculate each cell’s size-normalized acoustic scattering (SNACS), a parameter that increases monotonically with cell stiffness (*31*). SMR is three orders of magnitude faster than traditional AFM, and because it does not require that cells be replated, it circumvents artifacts arising from cellular adaptation to a second substrate. Our SMR measurements mirrored the results we had obtained by AFM, but with larger sample size, additional stiffness regimes, and enhanced statistical power (Fig. 1D). We conclude that B16F10 cells exhibit mechanoreciprocity, becoming stiffer in more rigid environments.

**Fig. 1:**
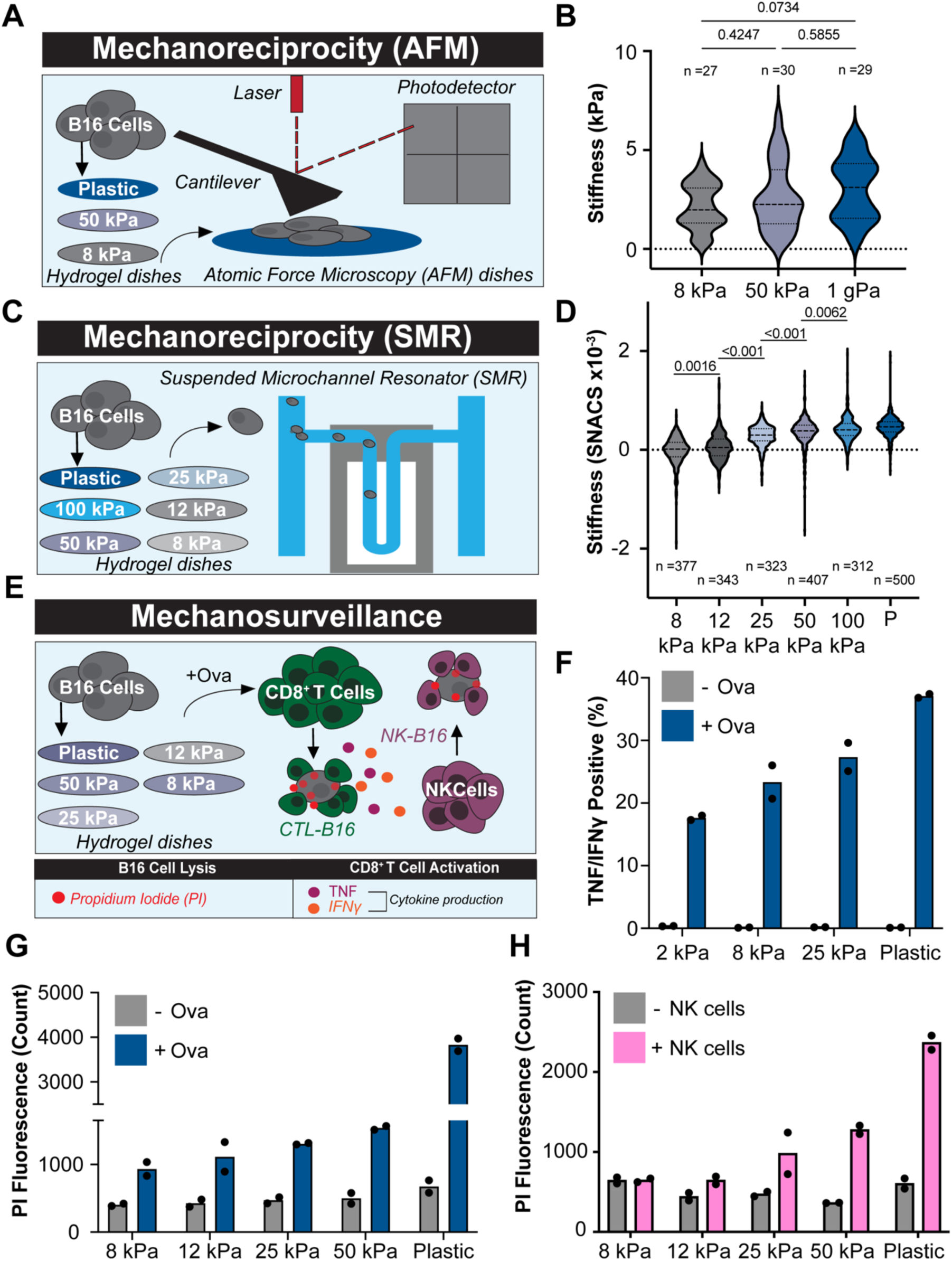
Substrate rigidity stiffens cancer cells and sensitizes them to cytotoxic lymphocytes. (A-D) B16F10 cells were cultured overnight on substrates of differential rigidity, and stiffness measurements of individual cells performed by AFM (A-B) or SMR (C-D). A and C show schematic diagrams of the approach. Note that the higher throughput SMR approach enabled assessment of more substrate rigidities. (B, D) Cell stiffness measurements at the indicated substrate rigidities, determined by AFM (B) and SMR (D). Violins encompass the entire data distribution, with dashed lines denoting the median and dotted lines indicating the upper and lower quartiles. Samples sizes (n=) are displayed above (B) or below (D) each violin. P-values calculated by one-way ANOVA. (E-H) B16F10 cells cultured overnight on substrates of differential rigidity were mixed with OT-1 CTLs in the presence of OVA, followed by quantification of B16F10 killing and CTL cytokine production. Analogous studies were performed using NK cells as cytotoxic lymphocytes. (E) Schematic diagram of the approach. (F) CTL cytokine production, expressed as the percentage of TNF^+^IFNγ^+^ CTLs after 5 h coculture with B16F10 cells. (G) B16F10 killing by CTLs, measured by propidium iodide uptake into dead cells after 5 h in the presence or absence of OVA. (H) B16F10 killing, measured by propidium iodide uptake after 5 h in the presence or absence of NK cells. All results are representative of at least two independent experiments.

Next, we examined the capacity of environmental stiffness to sensitize B16F10 targets to cytotoxic lymphocytes. B16F10 cells cultured on substrates of increasing rigidity were loaded with ovalbumin_257-264_ peptide (OVA) and then mixed with CTLs expressing the OT-1 T cell receptor (TCR), which recognizes OVA in the context of the class I major histocompatibility complex (MHC) protein H-2K^b^ (Fig. 1E). On stiffer substrates, B16F10 cells elicited markedly higher levels of CTL activation, which we measured by production of the inflammatory cytokines IFNγ and TNF (Fig. 1F). Target cell killing also increased with substrate rigidity (Fig. 1G), consistent with the interpretation that rigid environments render cancer cells more stimulatory to CTLs and therefore more vulnerable to their killing responses. B16F10 cells cultured on stiff substrates also became sensitized to killing by NK cells (Fig. 1H), indicating that this environmentally driven process affects multiple types of cytotoxic lymphocyte. Taken together, these results indicate that mechanoreciprocity can indeed promote the mechanosurveillance of cancer cells.

### Environmental rigidity sensitizes MTCs to cytotoxic lymphocytes *in vivo*

The capacity of substrate rigidity to dictate the stiffness and the immune vulnerability of cancer cells *in vitro*, taken together with prior studies showing that cytotoxic lymphocytes preferentially destroy stiffer target cells (*28-30*), prompted us to hypothesize that cancer cells invading stiff organs would be sensitized to immunosurveillance. To address this question, we intravenously (i.v.) injected Luciferase expressing (Luc^+^) B16F10 cells into immunocompetent C57BL/6 mice (wild type) or into perforin knockout (*Prf1^-/-^*) mice, which lacked the predominant pathway for cellular cytotoxicity (Fig. 2A). After three weeks, IVIS imaging of wild type recipients revealed frank metastases that were predominantly situated in the lungs (Fig. 2B). This distribution was consistent with previous work showing that i.v. injected B16F10 cells mainly seed through pulmonary circulation, rarely colonizing the bone (*32*). *Prf1^-/-^* animals exhibited a different outgrowth pattern; not only did metastases appear more quickly, but they also manifested in both the lungs and in the long bones of the leg (femurs) (Fig. 2B). The emergence of bone metastases was not simply the by-product of a generalized increase in outgrowth, as injection of 8-fold more B16F10 cells into wild type mice enhanced lung colonization without seeding detectable growth in the bone (fig. S1A). To quantify this change in metastatic site distribution, we determined the fraction of total IVIS signal coming from the legs (Fig. 2C and fig. S1B), and we also compared the change in femoral outgrowth to the change in total outgrowth (fig. S1C). Both approaches confirmed the selective expansion of femur metastasis in *Prf1^-/-^*mice.

**Fig. 2:**
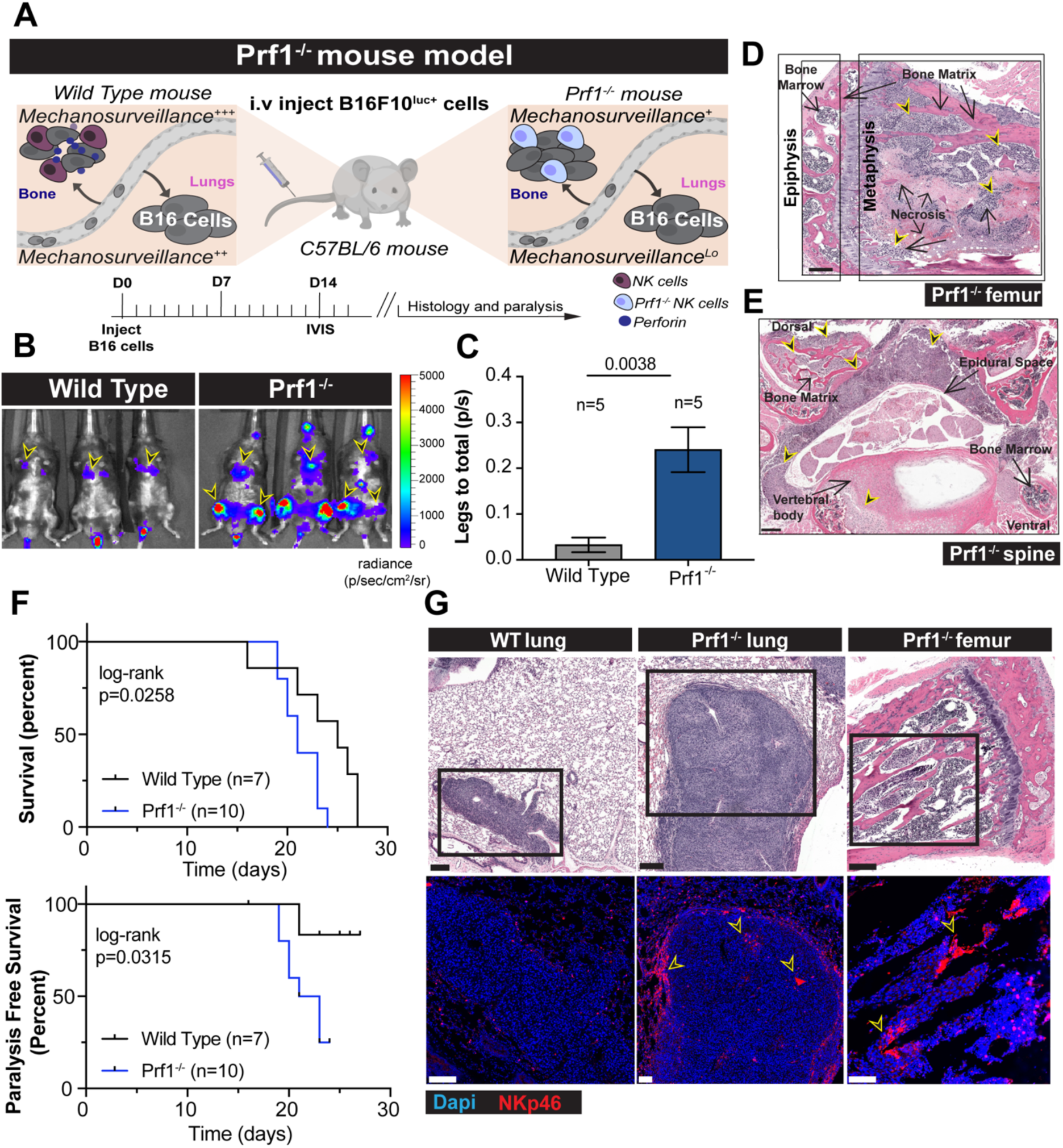
Preferential suppression of bone metastasis by cytotoxic lymphocytes. Luc^+^ B16F10 cells were injected i.v. into wild type and *Prf1^-/-^* mice, which were then monitored for metastatic colonization of the lungs and bone. (A) Schematic diagram of the experimental approach. (B) Representative IVIS images of tumor-bearing wild type and *Prf1^-/-^* mice, with metastatic burden in the lungs and femurs indicated by black and yellow arrowheads. (C) Quantification of relative femoral colonization, expressed as a ratio of IVIS signal in the legs to the total IVIS signal. Error bars denote standard error of the mean (SEM). Sample size is indicated above each bar. P-value calculated by unpaired Student’s t-test. (D-E) Representative H&E images of B16F10 bone metastases in the femur (D) and spine (E) of a *Prf1^-/-^*mouse. Epiphysis and metaphysis of the femur are indicated. Scale bars = 200 µm. Tumor cells indicated by black and yellow arrowheads. Scale bars = 200 μm. (F) Survival and paralysis-free survival of tumor-bearing wild type and *Prf1^-/-^* mice. (G) H&E images of representative metastatic tumors from the indicated tissues are shown above, with immunofluorescence staining of the boxed regions shown below, with NKp46^+^ NK cells in red. WT = wild type. Black and yellow arrowheads indicate NK cell clusters in the *Prf1^-/-^*bone. All results are representative of at least two independent experiments. Scale bars = 200 μm for H&E, 100 μm for immunofluorescence.

Histological analysis of femoral tumors in *Prf1^-/-^* mice highlighted the tendency of B16F10 cells to accumulate in the metaphyseal space beneath the epiphysial line, an area enriched in trabecular bone (Fig. 2D). Larger tumors occupied both the metaphysis and diaphysis (marrow shaft), suggesting initial metaphyseal colonization followed by expansion into the central femur. We also observed B16F10 metastases in the spines of *Prf1^-/-^*mice (Fig. 2E). These tumors occupied the vertebral body and/or articular processes, while also invading the epidural space. This growth pattern would be expected to compress the spinal cord, and consistently, we found that most *Prf1^-/-^* mice injected with B16F10 cells developed hind-leg paralysis, necessitating euthanasia (Fig. 2F). Conversely, little to no paralysis was observed in wild type recipients. The marked expansion of femoral and spinal colonization in *Prf1^-/-^* mice strongly suggested that cytotoxic lymphocytes are particularly important for restricting metastasis within the bone.

Immunofluorescence staining of B16F10 bone metastases from *Prf1^-/-^* mice revealed more pronounced accumulation of NK cells than CD8^+^ T cells (Fig. 2G and fig S1D), implying a more important role for the former in the surveillance of this cell line *in vivo*. Consistent with this interpretation, recipient mice treated with an NK cell depleting antibody exhibited enhanced metastatic colonization of the femur relative to mice receiving an isotype control antibody (Fig. 3A-C and fig. S2A-B). NK deficient animals, but not their isotype injected counterparts, also developed spine metastases and hind-leg paralysis after B16F10 injection (Fig. 3D-E), similar to our observations with *Prf1^-/-^* mice. By contrast, depletion of CD8^+^ T cells affected neither metastatic site distribution nor outgrowth (fig S2C-E). These results, which are consistent with prior data (*28, 33, 34*), indicate that the cytotoxic immunosurveillance of B16F10 metastases is predominantly mediated by NK cells.

**Fig. 3:**
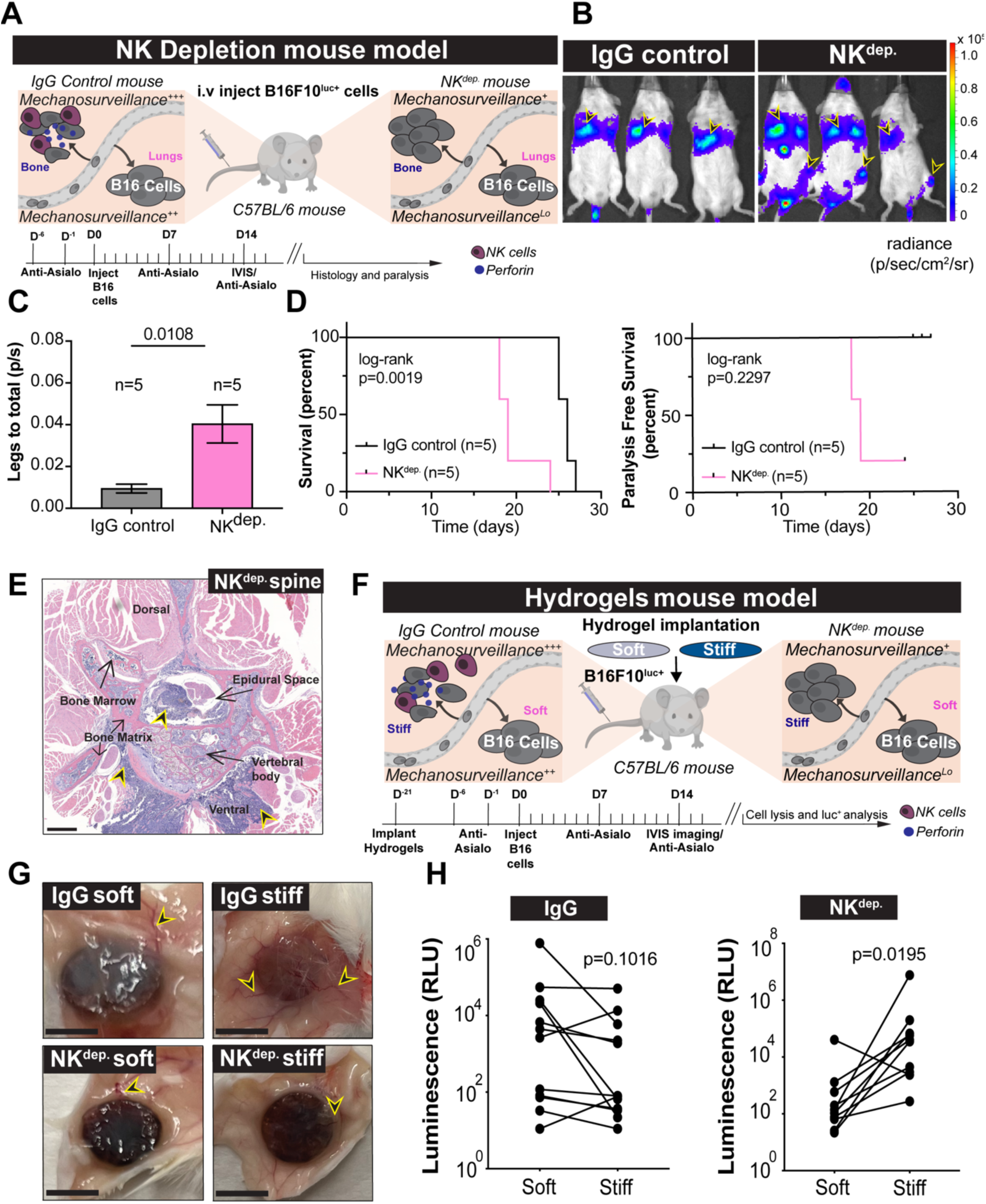
NK cells preferentially suppress metastatic outgrowth in rigid environments. Luc^+^ B16F10 cells were injected i.v. into IgG control and NK depleted (NK^dep.^) recipient mice, which were then monitored for metastatic colonization of the lungs and bone. (A) Schematic diagram of the experimental approach. (B) Representative IVIS images of tumor-bearing IgG control and NK^dep.^ mice, with metastatic burden in the lungs and femurs indicated by black and yellow arrowheads. (C) Quantification of relative femoral colonization, expressed as a ratio of IVIS signal in the legs to the total IVIS signal. Error bars denote SEM. Sample size is indicated above each bar. P-value calculated by unpaired Student’s t-test. (D) Survival and paralysis-free survival of tumor-bearing control and NK^dep.^ mice. (E) Representative H&E image of B16F10 bone metastasis in the spine of an NK^dep.^ mouse. Tumor cells indicated by black and yellow arrowheads. Scale bar = 500 μm. (F-H) C57BL/6 mice bearing stiff and soft subcutaneous hydrogel implants were treated with IgG control or NK cell depleting antibodies and then i.v. injected with Luc^+^ B16F10 cells. B16F10 colonization of the implants was assessed after 2 weeks. (F) Schematic diagram of the experimental approach. (G) Representative images of hydrogel implants three weeks after implantation, with black and yellow arrowheads denoting vascularization. Scale bars = 2.5 mm. (H) Implant colonization by B16F10 cells in IgG control (left) and NK^dep.^ (right) mice, measured by luciferase luminescence after hydrogel lysis. P-value calculated by paired Mann-Whitney test. All results are representative of at least two independent experiments.

To assess whether CD8^+^ T cells also preferentially target bone metastases *in vivo*, we performed analogous colonization experiments using EMT6, a triple negative breast cancer cell line. Compared to B16F10 cells, EMT6 cells express higher levels of class I MHC, rendering them more sensitive to CD8^+^ T cell-mediated lysis and less sensitive to NK cells (*35*). I.v. injection of EMT6 cells into wild type (BALB/c) recipients yielded predominantly lung metastases, with weak but detectable tumor formation in the femurs. Depletion of CD8^+^ T cells increased the colonization of both locations, as expected (fig. S3A). This increase was more pronounced in the femurs, however, consistent with stronger immunosurveillance in the bone microenvironment (fig. S3B-C). Hence, the capacity of cytotoxic lymphocytes to restrict metastatic outgrowth in the bone holds for different cancer types and lymphocyte subsets.

Although the experiments described above strongly suggest that rigid tissues, like the bone, promote cytotoxic lymphocyte immunosurveillance, they do not demonstrate that increased rigidity is sufficient to enhance anti-tumor immunity, independently of other environmental differences. To address this question, we turned to a synthetic implantable microenvironment system recently developed to study colonization by disseminated cancer cells (*36*). 6 mm × 1 mm hydrogel discs of differential rigidity (15wt% vs. 30wt% polyacrylamide hydrogel) were implanted subcutaneously in opposing flanks of C57BL/6 mice. Following a period of vascularization and engraftment, mice were treated with NK cell depleting antibodies or isotype control. Luc^+^ B16F10 cells were then i.v. injected and their relative colonization of soft (15wt%) and stiff (30wt%) hydrogel scaffolds was evaluated after three weeks by resection and Luciferase assay (Fig. 3F-G). Preferential colonization of the stiffer hydrogels was observed in 8 out of 9 NK-depleted mice (Fig. 3H), suggestive of an invasion and/or growth advantage for these cells in more rigid microenvironments. This predilection was completely absent in control mice (Fig. 3H), implying that cancer cells occupying more rigid hydrogel niches are also more sensitive to NK cell-mediated killing. The observation that environmental rigidification can, on its own, enhance the immune vulnerability of disseminated cancer cells *in vivo* provides a mechanobiological basis for the preferential targeting of bone metastases by cytotoxic lymphocytes.

### Immune pressure and the microenvironment jointly dictate the stiffness and gene expression of MTCs

The hypothesis that mechanoreciprocity within the metastatic microenvironment dictates the sensitivity of cancer cells to mechanosurveillance makes two crucial *in vivo* predictions. The first of these, addressed in the section above, is that cancer cells colonizing rigid microenvironments should be particularly vulnerable to cytotoxic lymphocytes. The second prediction is that MTCs in different organs should exhibit stiffness properties reflective of both environmental rigidity and the level of cytotoxic immune pressure. Specifically, MTCs colonizing rigid environments like the bone should be stiffer than cells from soft organs like the lung, due to mechanoreciprocity. Furthermore, MTCs from mice lacking cytotoxic lymphocyte activity should be stiffer than cells from immunocompetent animals, due to mechanosurveillance.

To evaluate these hypotheses, we i.v. injected wild type and *Prf1^-/-^*mice with GFP expressing (GFP^+^) B16F10 cells, and then used SMR to profile the stiffness of MTCs isolated from the lungs and femurs of *Prf1^-/-^* recipients and the lungs of wild type recipients (wild type mice do not develop bone metastases after i.v. injection) (Fig. 4A). To minimize contamination from surrounding tissue, viable GFP^+^CD45^-^ cells were FACS purified prior to SMR analysis. MTCs from *Prf1^-/-^* lungs were significantly stiffer than those derived from wild type animals (Fig. 4B), in line with the expectation that stiffer cancer cells are preferentially destroyed by mechanosurveillance. The corresponding MTCs from *Prf1^-/-^* femoral metastases were markedly stiffer than both lung samples (Fig. 4B), indicative of substantial mechanoreciprocity in the rigid bone microenvironment. To confirm and extend these findings, we performed analogous SMR analyses of MTCs derived from NK cell-depleted mice and isotype antibody-treated controls. The results of these experiments recapitulated the *Prf1^-/-^* studies, with MTCs from NK-depleted lung being stiffer than cells from isotype control lung, and cells from NK-depleted bone being stiffer than both lung samples (Fig. 4C). Thus, MTCs from biophysically distinct organs exhibit mechanical phenotypes consistent with the effects of both mechanoreciprocity and mechanosurveillance.

**Fig. 4:**
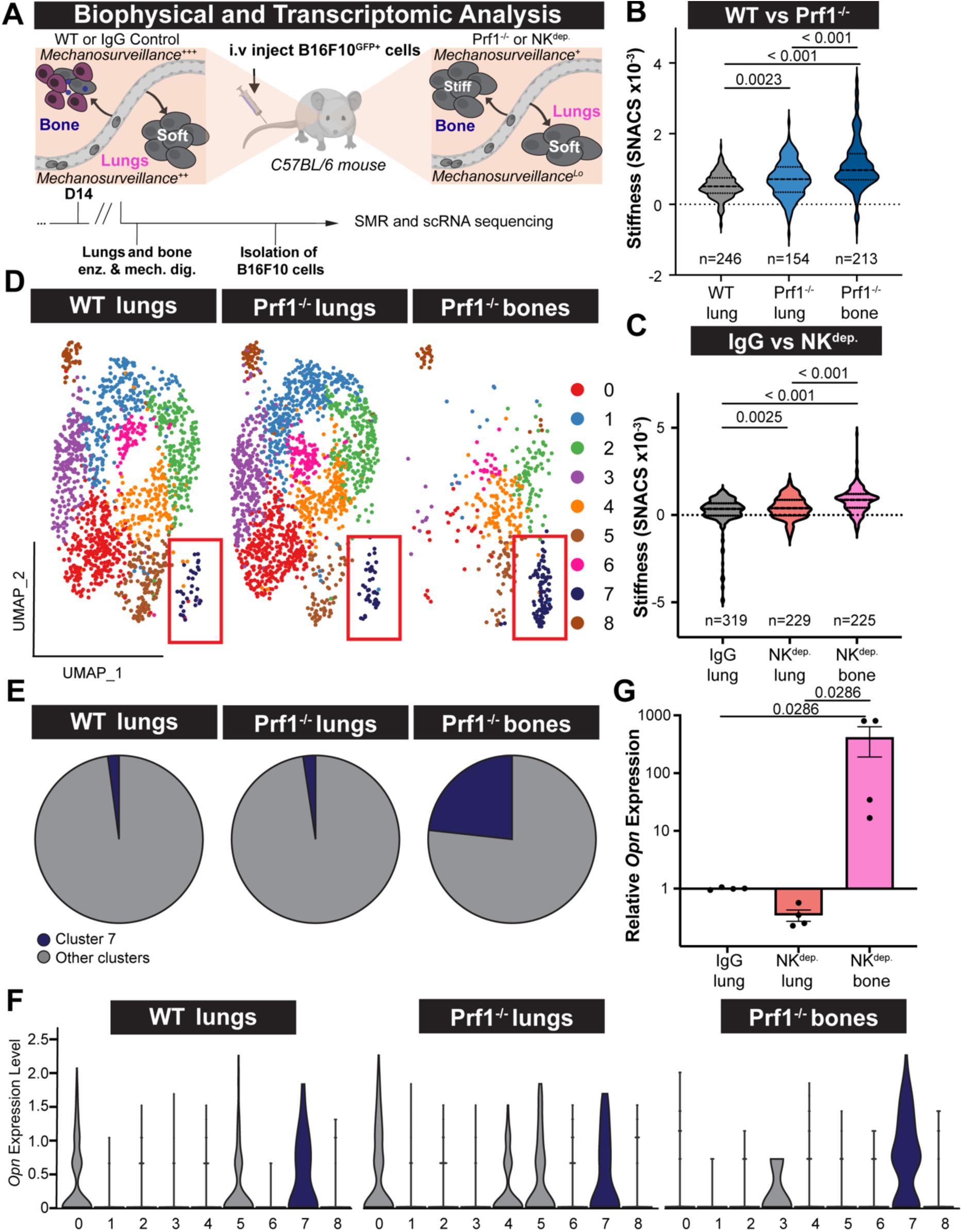
Environmental and immune regulation of MTC stiffness and gene expression. GFP^+^ B16F10 cells were i.v. injected either into wild type or *Prf1^-/-^* mice or into IgG control treated or NK cell depleted (NK^dep.^) mice. After 2 weeks, MTCs from the resulting metastases in the lungs of wild type and IgG control mice and the lungs and bones of *Prf1^-/-^* mice and NK^dep.^ mice were extracted and subjected to SMR and scRNA-seq. (A) Schematic diagram of the experimental approach. (B-C) SMR of B16F10 MTCs isolated from the indicated organs of wild type and *Prf1^-/-^*mice (B) or from IgG control and NK^dep.^ mice (C). Violins encompass the entire data distribution, with dashed lines denoting the median and dotted lines indicating the upper and lower quartiles. Samples sizes (n=) are displayed below each violin. P-values calculated by one-way ANOVA. (D) UMAP visualization of scRNA-seq data from the indicated lung and bone metastases. Cells are clustered based on transcriptional similarity and are color-coded by Seurat cluster identity. Cluster 7 has been boxed in each graph. (E) Pie charts showing the proportion of cluster 7 cells in each MTC sample. (F) Violin plot showing Opn expression levels across Seurat clusters in the indicated lung and bone metastases. Each violin represents the distribution of Opn expression within a cluster, with width indicating more cells. (G) qRT-PCR analysis of Opn expression levels in MTCs extracted from the indicated organs in IgG-treated versus NK^dep.^ tumor-bearing mice. Error bars denote SEM. P-values calculated by one-sample Wilcoxon test. All results are representative of at least two independent experiments.

To explore the molecular mechanisms underlying mechanoreciprocity and immune vulnerability *in vivo*, GFP^+^ B16F10 MTCs from the lungs of wild type mice and from the lungs and bones of *Prf1^-/-^* mice were subjected to single cell RNA-sequencing (scRNA-seq). Uniform manifold approximation and projection (UMAP) analysis of the resulting data revealed 9 distinguishable populations of cancer cells (Fig. 4D). Lung metastases from wild type and *Prf1^-/-^* animals contained all 9 populations, implying that cellular cytotoxicity does not drastically alter tumor composition in this organ. Nevertheless, subtle shifts in the size and composition of certain clusters were apparent, suggesting some degree of immune pressure. B16F10 composition differed dramatically in *Prf1^-/-^* bone, with several clusters shifting substantially in the UMAP plot or disappearing altogether. Most notable was a striking enrichment in cluster 7, which encompassed ∼25% of cancer cells in *Prf1^-/-^* bone (relative to ∼2% in the lung samples) (Fig. 4D-E).

Using differential gene expression analysis, we identified a suite of genes that were preferentially expressed in cluster 7 (fig. S4A), among them Osteopontin (*Opn*, also known as *Spp1*), which encodes a secreted glycoprotein and ECM component bound by several cell surface receptors, including CD44 and the α_v_β_3_ integrin (*37*). Opn was particularly intriguing because it is highly expressed by bone-resident cells and is known to promote osteoclast adhesion to bone matrix (*38, 39*). Furthermore, certain transformed cell types have been shown to upregulate Opn in response to substrate stiffness (*40, 41*) and upon infiltration of the bone microenvironment (*42*). Opn potentiates osteotropic metastases in mouse models through tumor cell adhesion and migration (*43-46*), while in human cancer patients, high Opn expression is both a feature of bone metastasis and an indicator of its prevalence (*47-51*). Using the Cancer Genome Atlas (TCGA), we documented both amplifications and deletions of Opn across a wide range of human cancers. Interestingly, amplification events are disproportionately represented in tumors from more rigid tissues (e.g. sarcoma), whereas Opn deletion is more prominent in diseases originating in softer organs (e.g. glioblastoma) (fig. S4B). Collectively, these observations suggest a role for Opn in cellular adaptation to rigid environments. In line with this hypothesis, we found that cluster 7 B16F10 cells from the bone expressed markedly higher levels of *Opn* than did cluster 7 cells in either lung sample (Fig. 4F). Similarly, qRT-PCR analysis revealed that bone-derived MTCs from NK-depleted mice expressed higher levels of *Opn* than did MTCs from isotype control and NK-depleted lungs (Fig. 4G). Taken together, these results document the selective expansion of a subset of B16F10 cells during bone metastasis and identify Opn as a molecular index of bone colonization.

### Opn controls cancer cell stiffness and mechanoreciprocity

The selective upregulation of *Opn* by metastatic B16F10 cells in the bone prompted us to interrogate its role as a potential mechanoregulator of metastatic site preference. To this end, we employed CRISPR/Cas9 to generate two independent Opn-knockout (KO) B16F10 cell lines, which we validated by qRT-PCR of the *Opn* transcript and sequence analysis of the *Opn* locus (fig. S5A-B). A control cell line was prepared in parallel using nontargeting (NT) guide (g)RNA. Opn-KO and NT control cells grew comparably *in vitro* (fig. S5C), indicating that Opn is dispensable for B16F10 proliferation. Both KO cell lines, however, were significantly less stiff than their NT counterparts, a phenotype that manifested using either SMR or AFM as the experimental readout (Fig. 5A-B). Opn-KO cells also contained significantly less F-actin (Fig. 5C-D), which was consistent with their enhanced deformability and also suggestive of a feedback relationship between Opn production and cytoskeletal strength. Notably, adding purified Opn protein to the culture medium restored the stiffness of Opn-KO cells to wild type levels (Fig. 5E). Furthermore, we found that an additional, “safe-harbor” control cell line, prepared using gRNA targeting the ROSA locus, retained wild type stiffness (fig. S5D). Hence, the mechanical softening of Opn-KO cells results specifically from loss of Opn rather than CRISPR-induced DNA damage.

**Fig. 5:**
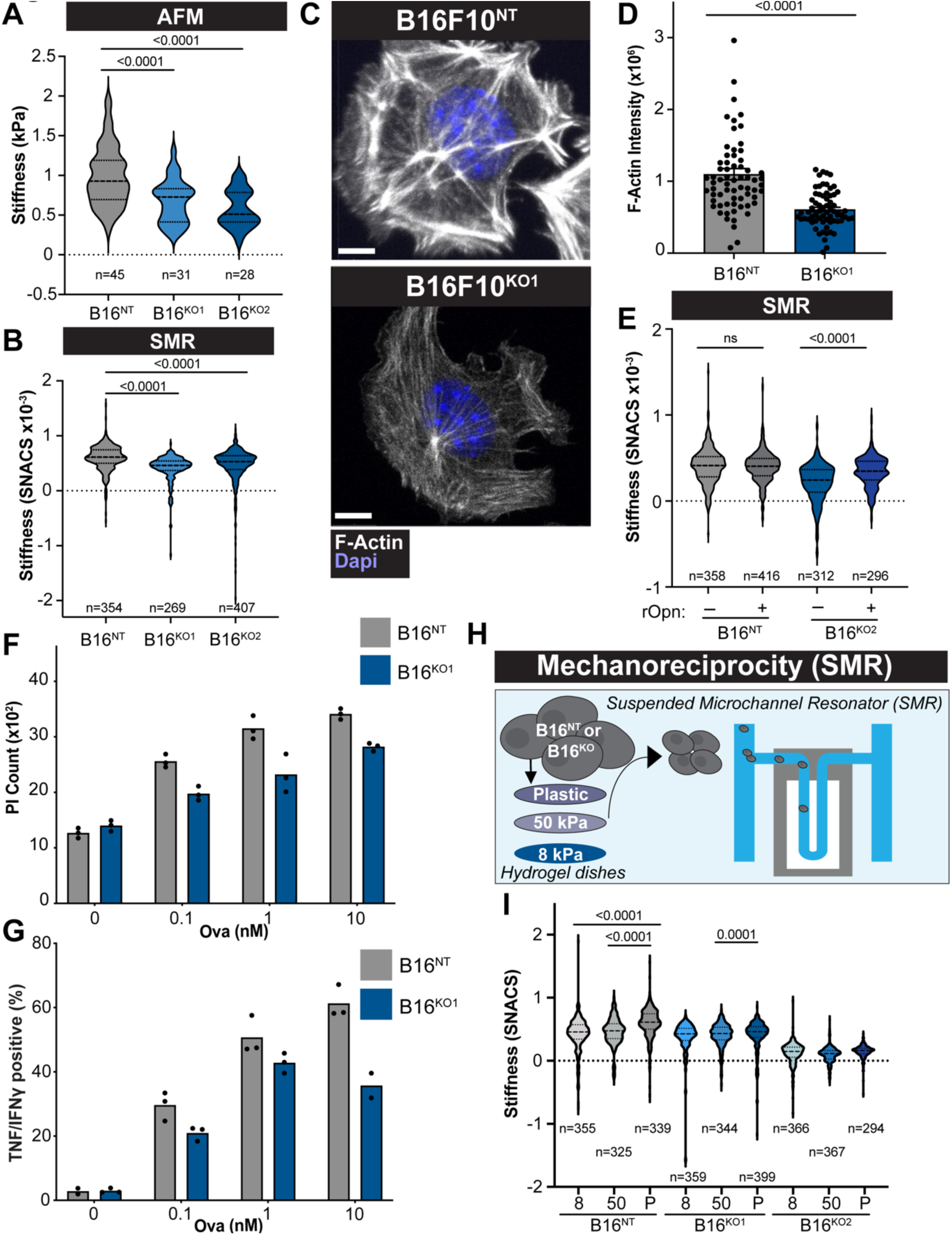
Opn mediates cancer cell mechanoreciprocity and cell stiffening. (A-C) The indicated Opn-KO B16F10 cell lines, along with nontargeting B16F10 control cells (NT), were subjected to AFM (A) and SMR (B) to measure their stiffness. (C-D) Phalloidin staining of Opn-KO1 and NT B16F10 cells. (C) Images of representative cells, with nuclei visualized by DAPI staining. Scale bars = 8 μm. (D) Quantification of F-actin intensity, with error bars indicating SEM. P-value calculated by unpaired Student’s t-test. (E) The indicated B16F10 cell lines were treated with purified Opn protein overnight and then subjected to SMR analysis. (F-G) The indicated B16F10 cells were loaded with increasing amounts of OVA and then mixed with OT-1 CTLs. (F) Target cell killing was measured by PI influx after 5 h. (G) CTL cytokine production, measured by intracellular staining for TNF and IFNγ, after 5 h. (H-I) NT or Opn-KO B16F10 cells were cultured overnight on substrates of differential rigidity and then subjected to SMR analysis. (H) Schematic diagram of the experimental approach. (I) Stiffness measurements at the indicated substrate rigidities. P = plastic. In A, B, E, and I, violins encompass the entire data distribution, with dashed lines denoting the median and dotted lines indicating the upper and lower quartiles. Samples sizes (n=) are displayed below each violin. P-values calculated by one-way ANOVA. All results are representative of at least two independent experiments.

We next determined whether Opn deficiency alters the sensitivity of B16F10 cells to cytotoxic lymphocytes. Both Opn-KO cell lines were less susceptible than control cells to CTL-mediated killing *in vitro* (Fig. 5F and fig. S6A), implying that Opn dependent stiffening renders B16F10 cells more stimulatory to CTLs. In line with this interpretation, CTLs cocultured with Opn-KO cells produced lower amounts of IFNγ and TNF (Fig. 5G and fig. S6B). The release of cytotoxic proteins, which we measured by surface exposure of the secretory lysosome marker Lamp1, was also impaired (fig. S6C). Notably, Opn-KO cells expressed normal amounts of class I MHC, indicating that their reduced stimulatory capacity was not caused by lower activating ligand expression (fig. S6D). Collectively, these results strongly suggest that Opn controls cancer cell mechanics and immune vulnerability.

Having found that Opn promotes cell stiffness, we next asked whether it might also influence mechanoreciprocity. To this end, we used SMR to measure the stiffness of Opn-KO and NT control B16F10 cells that had been plated on the aforementioned panel of substrates (Fig. 5H). NT controls exhibited mechanoreciprocity, as expected, increasing their stiffness in response to substrates of higher rigidity (Fig. 5I). By contrast, neither KO cell line exhibited any appreciable substrate-induced stiffening (Fig. 5I). These results indicate that Opn is crucial for B16F10 mechanoreciprocity and provide a potential mechanism by which this protein might facilitate adaptation to and colonization of rigid microenvironments like the bone.

To evaluate the importance of Opn for *in vivo* metastasis, we i.v. injected Luc^+^ Opn-KO or NT control B16F10 cells into wild type or *Prf1^-/-^* mice (Fig. 6A). As expected, NT control cells colonized the lungs of wild type recipients and both the lungs and the femurs of *Prf1^-/-^* animals (Fig. 6B-C, fig. S7A-C), indicative of a robust capacity for bone metastasis that is attenuated by cytotoxic lymphocytes. In stark contrast, Opn-KO cells were unable to colonize the bone in either group of recipients. The same pattern of results was observed using NK cell-depleted mice and isotype antibody-treated controls; NK-depleted mice readily developed NT B16F10 metastases in the bone but did not support detectable Opn-KO colonization in this niche (Fig. 6D-F, fig. S7D-F). Given that Opn-KO B16F10 cells proliferate comparably to their NT control counterparts (fig. S5C), these differences in bone colonization are unlikely to result from changes in post-seeding outgrowth. Rather, our results indicate that Opn, and potentially Opn dependent mechanoreciprocity, is required for effective infiltration of and biomechanical adaptation to rigid microenvironments. In line with this interpretation, comparative RNA-sequencing analysis of Opn-KO cells revealed decreased expression, relative to NT controls, of genes involved in extracellular matrix production, cytoskeletal remodeling, and adhesion (Fig. 6G-H). These results indicate that Opn-induced cell stiffening is coupled to a more adhesive and architecturally engaged cellular state, and they also suggest that adopting this state is necessary for bone colonization.

**Fig. 6:**
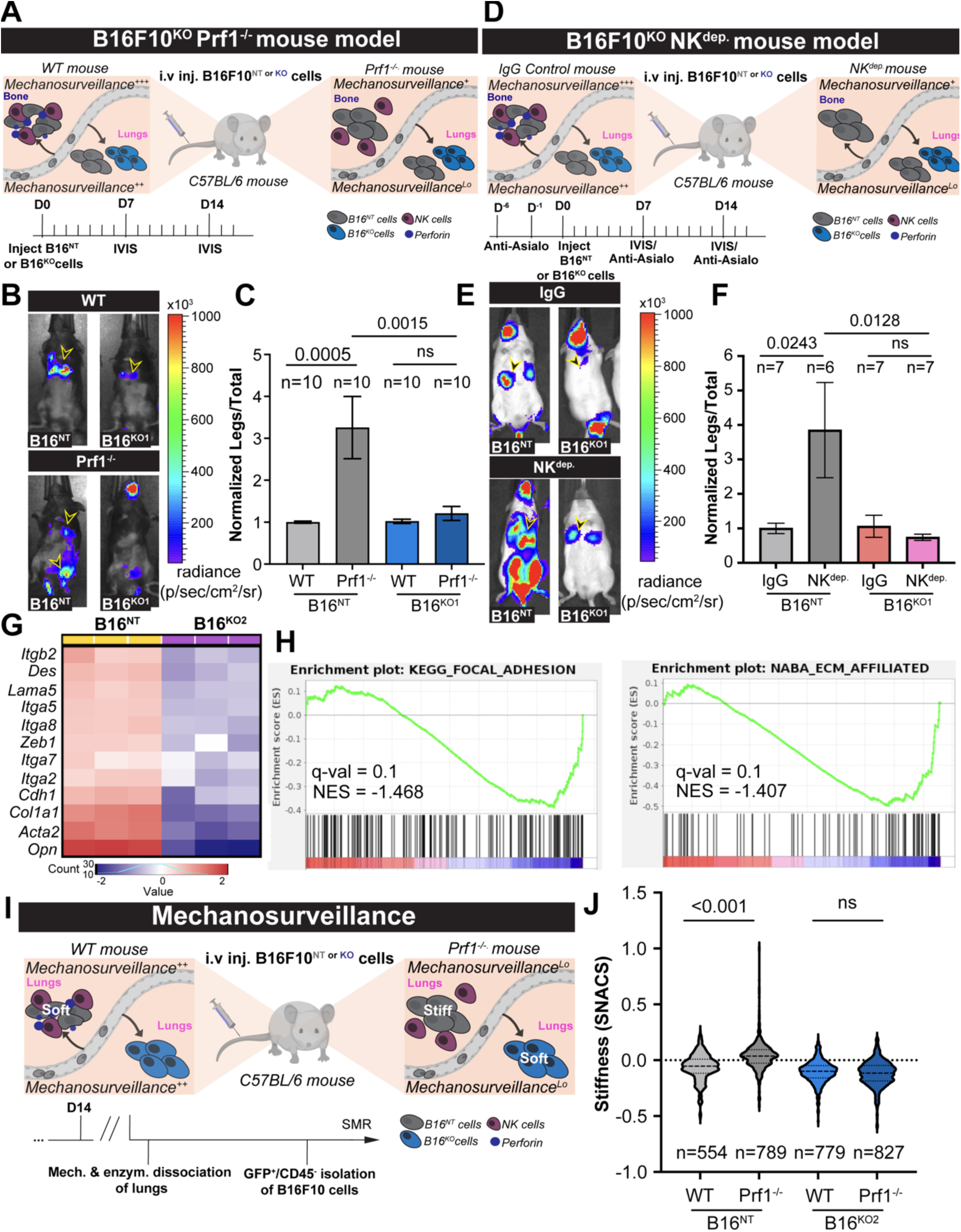
Opn is required for mechanoreciprocity and mechanosurveillance in vivo. (A-C) Luc^+^ NT or Opn-KO1 B16F10 cells were injected i.v. into wild type and *Prf1^-/-^* mice, which were then monitored for metastatic colonization of the lungs and bone. (A) Schematic diagram of the experimental approach. (B) Representative IVIS images of tumor-bearing wild type and *Prf1^-/-^* mice injected with the indicated B16F10 lines. (C) Quantification of relative femoral colonization, expressed as a ratio of IVIS signal in the legs to the total IVIS signal. (D-F) Luc^+^ NT or Opn-KO1 B16F10 cells were injected i.v. into IgG control and NK depleted (NK^dep.^) recipient mice, which were then monitored for metastatic colonization of the lungs and bone. (D) Schematic diagram of the experimental approach. (E) Representative IVIS images of tumor-bearing IgG control and NK^dep.^ mice injected with the indicated B16F10 lines. (F) Quantification of relative femoral colonization, expressed as a ratio of IVIS signal in the legs to the total IVIS signal. In B and E, metastatic burden in the lungs and femurs is denoted by black and yellow arrowheads. In C and F, error bars denote SEM, sample size is indicated above each bar, and P-values were calculated by one-way ANOVA. (G-I) NT and Opn-KO2 B16F10 cells were subjected to comparative bulk RNA-seq. (G) Heat map showing downregulation of selected ECM, adhesion, and cytoskeleton-related genes in Opn KO2 cells. (H) Gene Set Enrichment Analysis (GSEA) showing downregulation of genes related to focal adhesions (left) and ECM (right). NES = normalized enrichment score. (I-J) GFP^+^ NT or Opn-KO B16F10 cells were injected i.v. into wild type and *Prf1^-/-^* mice, and after 2 weeks, MTCs from the resulting lung metastases were extracted and subjected to SMR. (I) Schematic diagram of the experimental approach. (J) SMR of the indicated MTCs extracted from the indicated tumor-bearing mice. Violins encompass the entire data distribution, with dashed lines denoting the median and dotted lines indicating the upper and lower quartiles. Samples sizes (n=) are displayed below each violin. P-values calculated by one-way ANOVA. All results are representative of at least two independent experiments.

Finally, we investigated whether Opn influences mechanosurveillance *in vivo*. To this end, we applied SMR to mechanically profile GFP^+^ Opn-KO and NT control B16F10 MTCs extracted from the lungs of wild type and *Prf1^-/-^* mice (Fig. 6I). This approach was motivated in part by the observation that MTCs from *Prf1^-/-^* lungs expressed more *Opn* than the corresponding MTCs from WT mice (Fig. 4F), and would therefore be expected to elicit stronger cytotoxic lymphocyte activation. In line with our initial *in vivo* SMR experiments (Fig. 4B), NT MTCs from *Prf1^-/-^* animals were significantly stiffer than NT MTCs from wild type recipients (Fig. 6J). This result is consistent with the interpretation that stiffer MTCs are preferentially destroyed by patrolling cytotoxic lymphocytes. Remarkably, Opn-KO MTCs exhibited identical stiffness profiles regardless of whether they were purified from wild type or *Prf1^-/-^* mice (Fig. 6J). Hence, to the extent that Opn-KO cells experience immune selection *in vivo*, this selection is not based on their stiffness. Taken together with the experiments described above, these data identify Opn as a critical regulator of both the mechanoreciprocity and the mechanosurveillance of MTCs.

### The mechanical properties of human MTCs reflect environmental stiffness and immune pressure

The relevance of mechanosurveillance for anti-tumor immunity in clinical contexts remains largely unexplored. To address this issue, we examined publicly available scRNA-seq data sets that were generated to investigate patient responsiveness to ICB, an immunotherapeutic modality that unleashes tumor specific T cells, and would therefore be expected to enhance CTL-mediated mechanosurveillance. Our efforts focused on a large study comprising 41 breast cancer patients treated with one dose of pembrolizumab, which targets the inhibitory receptor PD-1 (*52*) (Fig. 7A). Biopsies were extracted just before pembrolizumab injection and 9-11 days later, providing a highly uniform assessment of early ICB-induced anti-tumor immunity in the tumor microenvironment. Importantly, patient responsiveness in this study was evaluated by the clonal expansion of T cells after pembrolizumab treatment, enabling us to infer the effects of specific cancer cell properties on T cell activation.

**Fig. 7:**
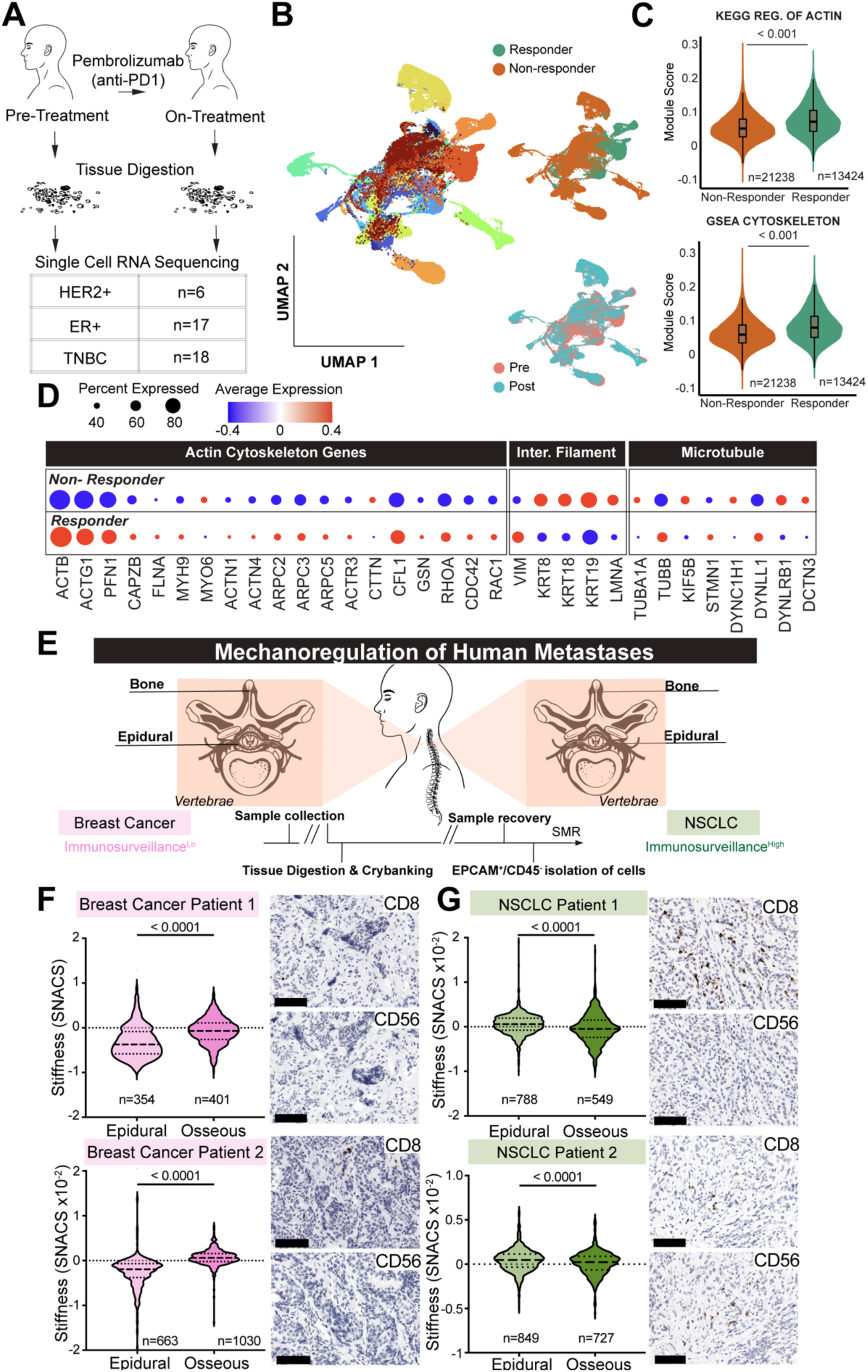
Cancer cell stiffness reflects environmental rigidity and immune pressure in human tumors. (A-D) scRNA-seq analysis of cytoskeletal gene expression in breast cancer patients treated with ICB. (A) Schematic diagram illustrating the study design: Breast cancer biopsies from patients with the indicated disease subtypes were collected before and 9-11 days after a single dose of anti-PD-1 immunotherapy, then subjected to scRNA-seq. (B) UMAP visualization of scRNA-seq data colored by patient ID (left), outcome (top right), and treatment status (bottom right). (C) F-actin cytoskeleton gene expression in non-responder versus responder tumors, calculated using data from pre-treatment samples. Module scores were generated using the KEGG “Regulation of Actin Cytoskeleton” pathway and a GSEA “Cytoskeleton” gene set. Embedded boxes indicate median and interquartile range. P-values calculated by Wilcoxon rank-sum test. (D) Dot plot showing relative levels of selected highly expressed components of the F-actin, intermediate filament, and microtubule cytoskeletons. (E-G) Comparative analysis of MTCs from the osseous and epidural regions of spinal metastases. (E) Schematic diagram illustrating the anticipated differences in the immunosurveillance of breast cancer versus NSCLC metastases. (F-G) Left panels, SMR of epidural versus osseous MTCs isolated from the spine of patients with metastatic breast cancer (F) or NSCLC (G). Right panels, representative IHC images showing CD8 and CD56 staining in epidural breast cancer (F) or NSCLC (G) sections. Scale bars = 100 μm. In F and G, violins encompass the entire data distribution, with dashed lines denoting the median and dotted lines indicating the upper and lower quartiles. Samples sizes (n=) are displayed below each violin. P-values calculated by unpaired Student’s t-test.

In UMAP plots, cancer cells were predominantly clustered by patient rather than by responsiveness to ICB (responder versus non-responder) or status of therapy (pre versus post) (Fig. 7B). This was not unexpected, as specific oncogenic drivers and underlying patient genetics were probably the predominant sources of variation among the tumors in this study. Next, we assessed the expression of cytoskeletal genes in all pre-treatment samples, with the goal of identifying predictive associations between cancer cell stiffness and ICB responsiveness. This analysis revealed a significant over-representation of gene sets encompassing actin and its regulators among patients that subsequently responded to ICB (Fig. 7C, fig. S8A). Over-representation of actin cytoskeletal genes in responders was particularly obvious for highly expressed proteins, such as actin itself (*ACTB* and *ACTG1*), factors that promote filament assembly (ARPC2, *CFL1*, and *PFN1*), bundling proteins (*ACTN1* and *ACTN4*), and small GTPases (*RHOA*, *RAC1*, and *CDC42*) (Fig. 7D). Highly expressed components of the microtubule and intermediate filament cytoskeletons did not exhibit this degree of uniform upregulation (Fig. 7D). The association between F-actin cytoskeletal genes and ICB-induced T cell activation in this patient cohort is consistent with the idea that cell stiffness potentiates lymphocyte-mediated anti-tumor responses via a mechanosurveillance mechanism. Interestingly, upon dividing the data set by breast cancer class, we found that the prognostic value of actin cytoskeletal gene expression applied only to triple negative breast cancer (TNBC), and not to estrogen receptor positive (ER^+^) or HER2^+^ tumors (fig. S8B). This result tracks with prior studies suggesting that TNBC is more responsive to ICB than other disease subtypes (*53, 54*).

By mapping the abundance of individual transcripts onto the UMAP space, we were able to discern notable differences in the expression of F-actin cytoskeletal genes (e.g. *ACTB*, *PFN1*) between cancer cell clusters, even between clusters from responder patients (fig. S8C). Similar cluster-to-cluster variation was observed for *B2M*, a major histocompatibility complex subunit that promotes T cell immunosurveillance by enhancing antigen presentation by cancer cells (*55*). Interestingly, certain responder cells with relatively low *B2M* expression exhibited high levels of *ACTB* and/or *PFN1* (fig. S8C), implying that, in some cases, mechanosurveillance can compensate for suboptimal antigen presentation in sensitizing cancer cells to immune attack.

Having identified indices of mechanosurveillance in patient samples, we next sought to interrogate the roles of environmental rigidity and mechanoreciprocity in human anti-tumor immunity. To this end, it was necessary to obtain patient samples that delineated the effects of the tumor microenvironment from those caused by patient-specific tumor etiology. Our approach centered on physically contiguous spinal metastases occupying both the osseous (bony) regions of vertebrae as well as the epidural space around the spinal cord, a softer environment containing fat, blood vessels, and connective tissue (Fig. 7E). By resecting samples from the osseous and epidural components of the same metastasis, we were able to compare isogenic MTC preparations derived from two mechanically distinct environments.

Paired osseous and epidural samples were obtained from two patients with breast cancer (one ER^+^HER2^-^, one ER^+^HER2^+^) and two patients with non-small cell lung cancer (NSCLC). No patient received ICB therapy prior to surgery. Samples were dissociated and FACS purified to isolate EpCAM^+^ MTCs, which were then applied to the SMR. Osseous MTCs from the breast cancer samples were substantially stiffer than their epidural counterparts (Fig. 7F), consistent with mechanoreciprocity in the bone microenvironment. Interestingly, this pattern was not observed in the NSCLC samples, where the osseous MTCs were actually a bit softer than MTCs from the adjacent epidural space (Fig. 7G). We reasoned that these discrepancies might reflect differential involvement of cytotoxic lymphocytes during metastatic colonization. NSCLC is considered to be more immunologically responsive than ER^+^ or HER2^+^ breast cancer, reflecting a higher mutational load (*56, 57*). Augmented mechanosurveillance by cytotoxic lymphocytes would be expected to preferentially deplete stiffer MTCs in the bone, thereby reversing the effects of mechanoreciprocity in this microenvironment.

To investigate this possibility, fixed sections from the same patient samples were stained for CD45 (a pan-leukocyte marker), CD3 (for T cells), CD8 (to identify CTLs), and CD56 (a marker for NK cells). Tumor domains were identified unambiguously by H&E staining of adjacent sections (fig. S8D). NSCLC metastases were characterized by marked lymphocytic infiltration of the tumor bed in both the osseous and epidural regions. In NSCLC patient #1, this phenotype was primarily driven by robust accumulation of CD8^+^ CTLs, whereas in NSCLC patient #2, roughly equivalent numbers of CD8^+^ and CD56^+^ cells were detected in the tumor bed (Fig. 7G). By contrast, the breast cancer metastases exhibited appreciably lower levels of lymphocyte infiltration. Few CD8^+^ and CD56^+^ lymphocytes were present in Breast cancer tumor #1, and although Breast cancer tumor #2 contained more CD8^+^ cells, almost all of them were segregated in interstitial regions between the lobule-like subdomains of the tumor bed (Fig. 7F). Interestingly, these interstitial domains contained clusters of large, CD45^+^ cells (fig. S8D), not unlike the macrophage-based lymphocyte exclusion barriers that have been observed in certain metastatic tumors (*58-60*). This structural feature, alongside the reduced lymphocyte infiltration seen in both breast cancer samples, may imply that these tumors were less subject to cytotoxic immunosurveillance than their NSCLC counterparts. Taken together with our SMR analysis of murine metastases, these results are consistent with the interpretation that human MTCs stiffen mechanoreciprocally in the rigid bone microenvironment, and then either persist in this state or are selectively eliminated in the face of robust cytotoxic lymphocyte mechanosurveillance.

## Discussion

In this study, we integrated biophysical assays, gene expression profiling, and metastatic models to dissect the role of environmental stiffness in shaping metastatic site distribution through differential immune vulnerability. We demonstrated that environmental rigidity influences metastatic outgrowth via the coupled effects of mechanoreciprocity and mechanosurveillance, and further identified Opn as a critical, cancer cell-derived mediator of both processes. Importantly, transcriptional, biophysical, and histological analysis of patient samples indicated that the mechanoregulatory paradigm we established in mice also applied to human cancer. We conclude that mechanoreciprocity plays a dual role during metastatic progression: facilitating the outgrowth of cancer cells in rigid tissues while concomitantly enhancing their sensitivity to cytotoxic lymphocytes (fig. S9). These results not only expand the scope of mechanobiology in metastasis but also provide a framework for exploring therapeutic interventions targeting the interplay between mechanical adaptation and immunosurveillance.

Our model positions cancer cells as mechanical interpreters, of sorts, translating environmental rigidity into a change in cell stiffness that lymphocytes can sense through the immune synapse. This intermediary role is consistent with our observation that Opn deletion abrogates B16F10 mechanoreciprocity while also markedly attenuating the capacity of stiff substrates to sensitize B16F10 cells to cytotoxic lymphocytes. It is important to note, however, that our data do not exclude the possibility that environmental rigidity may also modulate lymphocyte function via direct, extrasynaptic pathways. This would be in line with recent work indicating that the rigidity of a synthetic culture scaffold can influence antigen induced T cell expansion in a manner that is independent of antigen presenting cell stiffness (*61*).

It is now clear that tissues impose mechanical requirements for metastatic colonization (*9, 62*). However, whether these requirements are achieved via adaptation of cancer cells to their new environment or by selection of colonization competent cells within the larger metastatic pool remains a point of contention. We have found that B16F10 cells mechanically adjust to the rigidity of their substrate within 16 hours, before substantial proliferation or cell death has occurred. Furthermore, Opn-KO cells, which fail to mechanoreciprocate, also fail to colonize the bone *in vivo*. Taken together, these results strongly suggest that the mechanical adaptation of individual cancer cells to environmental rigidity is crucial for bone metastasis, at least in this experimental model. This interpretation does not rule out a role for selection in the colonization of stiff environments. It does, however, imply that if selection does contribute, it is not selection for cell stiffness *per se*, but rather for the capacity to stiffen in appropriate circumstances. Hence, mechanical adaptability itself emerges as a critical selectable trait, supported by gene products like Opn, which enable cancer cells to mechanoreciprocate. Similarly, our results are not inconsistent with mechanical memory (*40, 63, 64*), in particular the idea that MTCs retain biophysical characteristics imprinted by the primary tumor microenvironment, which influence their metastatic potential. They do suggest, however, that mechanical memory may not be encoded directly in cytoskeletal architecture, but rather in the capacity of that architecture to respond productively to environmental rigidity. The importance of prioritizing mechanical adaptability during metastasis, rather than a specific cellular mechanotype, makes sense given the variety of biophysically distinct environments MTCs must traverse during their dissemination.

Within the bone metaphysis, rigid trabecular bone matrix is closely juxtaposed with exceedingly soft bone marrow, generating a highly nonuniform mechanical environment. The fact that Opn, which drives stiffening on rigid substrates, is required for colonization of this niche strongly suggests that mechanoreciprocity to rigid, osseous components of this environment is an essential, early step in metastatic invasion. Subsequent expansion into marrow dominated spaces, however, would be expected to elicit a different kind of mechanoreciprocal response, one that enhances biomechanical variation within the metastatic tumor. Our scRNA-seq and SMR results are both consistent with this idea, revealing high levels of transcriptomic and mechanical diversity among bone MTCs. Mapping the mechanical and transcriptomic heterogeneity of MTCs onto the microarchitecture of bone environment will be an interesting topic for future research.

Opn has long been associated with bone metastasis in patients and in animal models (*42-51*). However, the precise manner through which it promotes colonization of this organ remains unresolved. Our results establish a biomechanical basis for Opn activity, demonstrating not only that it is upregulated in stiff environments, but also that it enables cancer cells to sense those environments. Given that Opn is a secreted protein that incorporates into the ECM, it is tempting to speculate that it functions as an “adhesion adaptor”, enabling cancer cells that express the α_v_β_3_ integrin or other Opn receptors to better adhere and respond to bone matrix. Such an adaptor role may be particularly critical under conditions of high bone mineralization, which was recently shown to attenuate integrin contacts (*65*). Opn-mediated adhesion could potentially trigger a positive feedback loop through which external stiffness prompts additional Opn secretion, which would further reshape the ECM to favor metastatic progression. Analogous feedback relationships have been shown to drive ECM remodeling and tumor progression in multiple contexts (*66*). Importantly, a role for secreted Opn as an adhesion adaptor is consistent with prior studies showing that Opn promotes osteoclast binding to bone matrix and that Opn accumulates at the interface between cancer cells and mineral bone in human metastases (*38, 39, 67*). This model raises the obvious question, however, of why cancer cells must express their own Opn, rather than exploit the presumably abundant Opn generated by other bone resident cells. It is possible that these cancer cell-extrinsic pools of Opn are bound by other cells or sequestered in microenvironments within the bone that are inaccessible or inhospitable to cancer cells for other reasons. Experiments targeting specific sources and isoforms of Opn will be necessary to address this issue.

Opn has been reported to inhibit CTL activation by engaging CD44, a cell surface receptor expressed by both T cells and NK cells (*68, 69*). This immunosuppressive mechanism contrasts with our results indicating that Opn mechanically sensitizes MTCs to cytotoxic lymphocytes. Whether the activating or the inhibitory process predominates likely depends on contextual factors such as environmental rigidity, the abundance of Opn, and whether that Opn is accessible to cancer cells, T cells, or both. Regardless, the capacity of Opn to both promote and antagonize cytotoxic lymphocyte function could potentially complicate efforts to target the protein therapeutically. While blocking both the immunosuppressive and mechanoreciprocal activities of Opn would presumably be desirable to prevent initial MTC infiltration of the bone, enhancing Opn dependent mechanoreciprocity while preventing CD44-based T cell suppression may be a better way to combat bone metastasis once it is established. Mechanistically delineating the immunosuppressive and mechanoregulatory activities of Opn could enable investigators to leverage the immunostimulatory benefits of this protein while simultaneously avoiding lymphocyte suppression.

Targetable cancer vulnerabilities canonically exhibit two key properties: first, they are required for disease progression, and second, they are specific to cancer cells. Mechanoreciprocal stiffening in rigid microenvironments satisfies the first criterion because it is a necessary, unavoidable step in metastatic progression in organs like the bone. This behavior is not unique to cancer cells, however, raising the question of how it serves as a specific trigger for anti-tumor immunosurveillance. The answer to this question likely lies in the combinatorial nature of immune recognition. In metastatic cells, mechanoreciprocal stiffening occurs in the context of a permissive constellation of surface ligands that is presumably not present in other bone resident cell types. This allows an otherwise normal biophysical feature to become a differential index of dysregulation that triggers cytotoxic lymphocyte attack. By combining mechanosensing with multimodal molecular recognition, immune cells expand both the scope and the specificity of their surveillance function.

## Acknowledgements

We thank Z. Eraslan and J. Zippin for assistance with CRISPR/Cas9-base deletion; P. Manoj, A. Shoenfeld, C. Rudin, N. Socci, X. Xiang, J. Wolchok, and T. Merghoub for advice with patient data analysis; the MSKCC Flow Cytometry and Molecular Cytology Core Facilities for assistance with FACS and imaging, respectively; T. Tammela, A. Boire, and E. Lee for critical reading of the manuscript; and members of the M. Huse lab for advice and encouragement.

## Authors contributions

Y. E., M. T.-L., A. H., Y. Z., B. Y. W., O. B., S. R. M., and M. H. designed the experiments. Y. E., M. T.-L., A. H., Y. Z., Z. W., A. Y., M. D., S. V., Y. H. K., and T. A. B. collected the data. Y. E., M. T.-L., A. H., Y. Z., Y. H. K., T. A. B., K. K. H. Y., O. B., and M. H. analyzed the data. Z. W., J.-G. K., and J. L. provided critical reagents. J. M., J. L., S. R. M., and M. H. provided research support. Y. E. and M. H. wrote the paper with input from A. H., Z. W., K. K. H. Y., J. M., J. L., O. B., and S. R. M.

## Supplementary Materials

### Materials and Methods

#### Cells and cell culture

B16F10 cells were cultured at 37 °C in RPMI supplemented with 10% FBS, 1 mM sodium pyruvate, 2 mM L-glutamine, 50 U/ml penicillin, and 50 μg/ml streptomycin. EMT6 cells were cultured at 37 °C in Waymouth medium with 10% FBS, 1 mM sodium pyruvate, 2 mM L-glutamine, 50 U/ml penicillin, 2 mM GlutaMAX, and 50 μg/ml streptomycin. HEK293T cells (used to generate ecotropic retrovirus) were cultured at 37 °C in DMEM supplemented with 10% FBS, 1 mM sodium pyruvate, 2 mM L-glutamine, 50 U/ml penicillin, and 50 μg/ml streptomycin. To generate OT-1 CTLs, splenocytes from OT-1 αβTCR transgenic mice were mixed with congenic splenocytes pulsed with 100 nM OVA and cultured in RPMI medium with 10% FBS and 0.55 mM β-mercaptoethanol. After 24 hours, cells were supplemented with 30 IU/ml IL-2 (NIH BRB Repository) and split as needed. Functional assays were performed after 7 days in culture. Murine NK cells were isolated from C57BL/6J splenocytes using negative selection (NK cell isolation kit, MACS, 130-115-818) and incubated overnight with 1000 U/ml IL-2. B16F10 cells were genetically modified to generate Opn and Rosa knockout (KO) lines using CRISPR/Cas9. Guide RNAs (gRNAs) targeting the *Opn* and *Rosa* loci were designed and synthesized by Synthego. Cells were transfected with multiguide gRNAs and Cas9 using Lipofectamine, followed by clonal expansion. Successful knockouts were confirmed by Sanger sequencing to validate genomic edits and by qRT-PCR to assess transcript depletion. Recombinant Opn (R-OPN) was used to rescue phenotypic effects in Opn KO cells. Cells were treated with R-OPN at 1 μg/ml for 24 hours before functional assays.

#### Animal Studies

The animal protocols for this study were approved by the Institutional Animal Care and Use Committee of Memorial Sloan Kettering Cancer Center. For B16F10 metastasis assays, 4-6 week old male wild type mice (C57BL/6J (Strain #00064) or B6(Cg)-*Tyr^c-2J^*/J (Strain #00058)) and CByJ.B6-Prf1tm1Sdz/J (Strain #007079) (*Prf1^-/-^*) mice were purchased from Jackson Laboratory. Mice were randomly assigned to groups for *in vivo* experiments. For EMT6 metastasis assays, 4-6 week old female BALB/c mice were purchased from Jackson Laboratory. OT-1 CTLs for *in vitro* assays were generated from 2-6 month old male and female OT-1 αβTCR transgenic mice, and NK cells were isolated from 2-4 month old male and female C57BL/6J mice, both obtained from Jackson Laboratory. All animals were housed under specific pathogen-free conditions.

#### Human Studies

Spinal metastases containing both epidural and osseous components were collected from breast and lung cancer patients indicated for surgery. Human tissues were obtained under Memorial Sloan Kettering Cancer Center (MSKCC) Institutional Review Board-approved protocol 17-593, titled “Defining the Immunological Tumor Microenvironment on Metastatic and Primary Spine Tumors.” Clinical information was abstracted from medical records and de-identified. Informed consent was obtained from all patients.

#### Metastasis Assays

For B16F10 metastasis assays, unless otherwise stated, 4 × 10⁵ cancer cells were injected into the tail vein of 4-6 week old C57BL/6J mice, and 2 × 10⁵ cancer cells into the tail vein of 4-6 week old *Prf1^-/-^* mice. Mouse hair was removed using clippers to prevent interference with bioluminescent imaging (BLI). Metastatic burden in the lungs and femurs was quantified weekly following retro-orbital injection of D-luciferin (150 mg/kg) and imaging using the IVIS Spectrum Xenogen instrument (Caliper Life Sciences) equipped with Living Image software v.2.50. For NK depletion experiments, 4 × 10⁵ (IgG control) or 2 × 10⁵ (NK depletion) B16F10 cells were injected into the tail vein of 4-6 week old B6(Cg)-Tyrc-2J/J mice (Jackson Labs, 000058). NK cell depletion was performed by intraperitoneal (i.p.) injection of anti-asialo GM1 antibody (Wako Chemicals, 986-10001), as previously described (*70*), 6 days and 1 day before tail vein injection of cancer cells and once weekly thereafter. CD8⁺ T cell depletion was achieved by administering 250 μg of InVivoMab anti-mouse CD8α antibody (clone 53-6.7, BioXCell, BE0004-1) or IgG2a control (BioXCell, BE0089) by i.p. injection 2 days and 1 day before tumor delivery, followed by weekly injections, as previously described (*28*). Survival and paralysis-free survival (defined as the time until bilateral hind leg paralysis) were tracked for all mice. Mice were euthanized upon reaching the endpoint criteria of paralysis, metastatic burden in the lungs, or other signs of significant distress.

#### Killing, degranulation, and cytokine production assays

For CTL functional assays, cancer cell targets were cultured overnight on fibronectin-coated 96-well plates in the presence of 20 ng/mL IFNγ (to enhance class I MHC expression), loaded with varying concentrations of OVA for 2 hours, and washed three times with medium. To assess killing, OT-1 CTLs were added at a 4:1 effector-to-target (E:T) ratio, and the lysis of target cells (GFP^+^CD8^−^) was tracked using the Incucyte S3 (Sartorius). PI (Thermo Fisher Scientific) was used to mark dead cells, and images were taken every hour for 8 hours to capture the kinetics of cell death. To assess lytic granule secretion, CTLs were mixed with B16F10 targets at a 2:1 E:T ratio and incubated for 90 minutes at 37 °C in the presence of eFluor660-conjugated anti-Lamp1 antibody (1 μg/ml, Clone 1D4B, eBiosciences). Cells were stained with anti-CD8a antibody, and the percentage of Lamp1^+^ CTLs (CD8^+^) was quantified by flow cytometry. For cytokine production, CTLs were added at a 2:1 E:T ratio and incubated for 4 hours at 37 °C in the presence of BD GolgiPlug protein transport inhibitor (BD Biosciences). Cells were then stained with anti-CD8a, fixed, permeabilized using the BD Cytofix/Cytoperm kit, and labeled with PE-conjugated anti-TNF (BioLegend, 506306) and PE/Cy7-conjugated anti-IFNγ (BioLegend, 505826) antibodies. The percentage of cytokine-producing CTLs (CD8^+^) was analyzed by flow cytometry. For NK cell functional assays, cancer cell targets were cultured overnight on fibronectin-coated 96-well plates, mixed with NK cells at a 4:1 ratio, and target cell lysis was measured using the Incucyte S3. PI flux was used to quantify cell death, with images taken every hour for 8 hours to track real-time killing. For assays using hydrogel substrates, hydrogel arrays (Matrigen) were coated with 10 µg/mL fibronectin (FN, Millipore Sigma) in PBS at 37°C for 2 hours to allow sufficient protein coating. Following incubation, excess FN was aspirated, and the wells were washed once with PBS to remove any unbound FN. B16F10 cells were then plated at a density of 30,000 cells per well in complete growth medium and incubated at 37°C overnight. Subsequent cocultures were performed as described above.

#### *In vitro* cell growth and proliferation assays

To assess proliferation, B16F10 cells were labeled with CellTrace Violet (CTV, Thermo Fisher) as per the manufacturer’s protocol, and CTV dilution was measured daily by flow cytometry to monitor cell division.

#### Histology

Murine lung and bone tissue was perfused and fixed overnight at 4°C with 4% PFA. Fixed bone was then decalcified in 10% EDTA (pH 7.4) at 4°C for 2-3 weeks with periodic monitoring to ensure complete decalcification. The tissues were then dehydrated through a graded ethanol series, cleared in xylene, and embedded in paraffin. Sections were stained for NKp46 (R&D Systems, Cat # AF2225) and CD8 (Cell Signaling Technology, Cat # 98941). Imaging was performed using a Pannoramic Scanner fitted with a 20×/0.8NA objective (3D Histech), and the results visualized using CaseViewer software (3D Histech).

#### Atomic Force Microscopy (AFM)

Cells were seeded on glass-bottom Petri dishes (FluoroDish FD5040) coated with FN and maintained in complete RPMI medium supplemented with 10 mM HEPES pH 7.0 during the acquisition of stiffness maps. Experiments were conducted at 37 °C using an MFP-3D-BIO AFM microscope (Oxford Instruments) with cantilevers fitted with 5 μm diameter colloidal borosilicate probes (nominal spring constant k = 0.1 N/m, Novascan). The exact spring constant of the cantilever was determined before each experiment using the thermal noise method, and its optical sensitivity was calibrated using a PBS-filled glass-bottom Petri dish as an infinitely stiff surface. Each session involved testing 10-12 cells from each experimental group. Bright field images of each cell were captured during AFM measurements using an inverted optical objective (Zeiss AxioObserver Z1) integrated with the AFM system. Stiffness maps of 60 μm × 60 μm (18 × 18 points) were acquired in areas containing both cells and substrate at a rate of 1.5 Hz for a single approach/withdrawal cycle. A trigger point of 1 nN was set to ensure a penetration depth of 1-2 μm. Force curves for each map were fitted using the Hertz model (Igor Pro, Wavemetrics). Data fitting was performed within the first 1 μm of indentation, specifically in the range of 0 to 50% of the maximum applied force. The following settings were used for the fitting: tip Poisson’s ratio ν_tip = 0.19, tip Young’s modulus E_tip = 68 GPa, and sample Poisson’s ratio ν_sample = 0.45. Stiffness histograms were obtained by identifying the stiffness values associated with each individual cell (excluding the substrate values). Data from each sample were pooled as a single population. Measurements made < 500 nm above the substrate were excluded from the analysis.

#### Suspended Microchannel Resonator (SMR)

Single-cell size-normalized acoustic scattering (SNACS) was measured using a previously described SMR-based method (*31*). Before each set of measurements, the SMR was treated with 0.25% Trypsin-EDTA for 30 minutes, followed by a 3-minute wash with 10% bleach and a final 5-minute rinse with DI-H_2_O. After cleaning, the SMR was passivated with 1 mg/mL poly(L-lysine)-graft-poly(ethylene glycol) (PLL-g-PEG) in H_2_O for 10 minutes at room temperature, followed by a 5-minute rinse with PBS + 2%FBS (FACS buffer). All measurements were performed at room temperature in FACS buffer. The SMR was briefly rinsed with the corresponding buffer between each experiment. During the measurements, cells were loaded into the SMR through 0.005-inch inner diameter fluorinated ethylene propylene (FEP) tubing. Fluid flow across the SMR was controlled using three independent electronic pressure regulators (MPV1, Proportion Air) and three solenoid valves (S070, SMC). A consistent differential pressure was applied across the SMR to maintain constant shear forces and data acquisition rate during cell measurement. All regulators, valves and data acquisition systems were operated by custom software developed in LabVIEW 2017 (National Instruments). The vibration frequency of the SMR cantilever was continuously monitored during measurements. Frequency shifts were used to quantify the node deviation (ND), which reflects acoustic scattering of individual cells, and the buoyant mass (BM) of the cells. ND is size-dependent, so it must be normalized to enable comparison across cells of varying volumes. We applied a normalization method as described in (*31*). To determine cell volume (*V*), we first we first measured the average cell volume 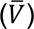 using a Coulter Counter (Beckman Coulter). Individual cell volumes were then inferred using the relationship:

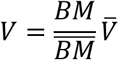

where 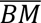 is the mean buoyant mass, estimated by fitting the distribution of buoyant mass measurements to a log-normal function. We then computed the size-normalized acoustic scattering (SNACS) value for each cell using the equation:

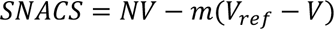

Here, *NV* = *ND*/*V* is the volume-normalized node deviation, and *m* is the slope obtained from a linear regression of NV versus V across the population. The reference volume *V_ref_* was defined as the median cell volume.

#### Immunofluorescence imaging and quantification

B16F10 cells were fixed in 2% PFA, washed in PBS, and then labeled with Alexa Fluor 647-labeled phalloidin (1:400, Invitrogen) and DAPI (1:1000, Sigma). To quantify F-actin intensity, phalloidin images was subjected to intensity thresholding in Imaris (Bitplane) to establish the space occupied by cells, after which the average intensity of phalloidin within the cellular volume was determined.

#### Hydrogel Scaffold Fabrication

Hydrogel scaffolds were fabricated following previously reported methods (*71*). Soda lime glass beads were sorted using an Advantech Sonic Sifter to ensure a consistent size range, with ∼8% deviation. The beads were dispersed in deionized water and gradually loaded into an 8 × 35 mm glass vial to a height of 2–2.5 mm. They were then mechanically packed into a lattice structure using an ultrasonic water bath. Preparations were then dried in a 60°C oven and thermally annealed for 4 hours in a furnace at temperatures between 650°C and 680°C, depending on the bead size. A hydrogel precursor solution was prepared immediately before use, consisting of 15% (soft) or 30% (stiff) acrylamide monomer, 1.5 wt% bis-acrylamide crosslinker, 0.2 vol% N,N,N′,N′-tetramethylethylenediamine accelerator, and 0.2 vol% 2-hydroxy-2-methylpropiophenone photoinitiator in nitrogen-purged deionized water. The precursor solution (150 µL) was infiltrated into the glass bead template and centrifuged at 4,000 × g for 15 minutes. It was then polymerized under a 15 W ultraviolet light source for 15 minutes. The polyacrylamide hydrogel–glass templates were removed from the glass vials the following day to ensure complete polymerization. Any excess hydrogel was removed by scraping the glass bead template with a razor blade on all surfaces. Glass beads were selectively dissolved in alternating washes of an acid solution: a 1:5 dilution of hydrofluoric acid in 1.2 M hydrochloric acid and 2.4 M hydrochloric acid. The washes were performed on a shake plate, with solutions being changed every 4 hours until the beads were fully dissolved. Scaffolds were thoroughly washed with deionized water to remove any residual acid and then lyophilized. After lyophilization, the scaffolds were resuspended in Cryomatrix embedding resin and cut to a thickness of 1 mm using a CryoStar NX70. Following further washing in deionized water, the scaffolds were sterilized with 70% ethanol and stored at 4°C in sterile phosphate-buffered saline (PBS) solution. The final pore dimension of the optimized scaffolds used in the study was 300 ± 16 µm.

#### Subcutaneous Hydrogel Implantation

Mice were anesthetized using 1.5% isoflurane, and their dorsal fur was removed with electric clippers followed by Nair hair removal cream. The skin was sterilized using 70% isopropyl alcohol prep wipes and povidone-iodine to minimize the risk of infection. Prior to the surgical procedure, each mouse received a subcutaneous injection of meloxicam (2 mg/kg) for analgesia. Two small horizontal incisions (2 mm) were made in the left and right flanks of the mouse. A subcutaneous pocket was carefully created at each incision by inserting surgical scissors and gently expanding them. Each pocket was then implanted with a single hydrogel scaffold. One scaffold was composed of 15% polyacrylamide, and the other scaffold was composed of 30% polyacrylamide, allowing for a direct comparison between the two materials within the same animal. The incisions were closed using two Reflex 7-mm wound clips, ensuring secure closure and promoting optimal healing. Postoperative care included daily meloxicam treatment for 3 days to provide continuous pain relief. The wound clips were removed after 7 days to allow for full tissue recovery.

#### Mouse Tissue Collection and Processing

All plasticware used in the procedure was precoated with 0.05% BSA, and centrifuges were pre-cooled to 4°C. Lungs and knees were digested in 1 mL of digestion buffer (2 mL Collagenase I [7.5 mg/mL], 1.49 mg DNase I, 8 mL 1% BSA in HBSS++) per mouse. For lung tissue, tumors were resected and digested in 10 mL of digestion buffer for 20 minutes at 37°C with rotation. After digestion, the material was strained, washed with 5 mL of SMEM + 2% FBS, and resuspended in 1 mL of SMEM + 2% FBS. Red blood cells were lysed with 10 mL of 1× Pharm Lysis for 1 minute, neutralized with 10 mL of SMEM + 2% FBS, and strained again. Cells were Fc-blocked (1:200) for 10 minutes at 4°C, followed by staining with anti-CD45-PerCPCy5 (1:200) and DAPI. Finally, cells were resuspended in 1 mL of PBS + 2% FBS for sorting. For knee tissue, the bones were cleaned, crushed in 1 mL of digestion buffer, and transferred to precoated Eppendorf tubes. An additional 1 mL of digestion buffer was added, and the samples were incubated at 37 °C for 30 minutes, with gentle pipetting every 10 minutes to facilitate digestion. Cells were processed following the same procedure as the lung samples. Both lung and knee tissue samples were subjected to FACS on a BD FACSAria to isolate GFP^+^CD45^-^DAPI^-^ cancer cells.

#### Human Tumor Tissue Processing

Tumors isolated from the spine were digested in 5 mL of digestion buffer (2 mL Collagenase I [7.5 mg/mL], 1.49 mg DNase I, 8 mL 1% BSA in HBSS++) for 20 minutes at 37°C with rotation. After digestion, the material was strained, washed with 5 mL of SMEM + 2% FBS, and resuspended in 1 mL of SMEM + 2% FBS. Red blood cells were lysed with 10 mL of 1× Pharm Lysis for 1 minute, neutralized with 10 mL of SMEM + 2% FBS, and strained again. Cells were Fc-blocked (1:200) for 10 minutes at 4°C, then stained with anti-CD45-FITC (1:200) and anti-EPCAM-eFluor660 (1:200), followed by DAPI staining. Cells were then resuspended in 1 mL of 2% FBS PBS for sorting. The human tumor samples were subjected to FACS on a BD FACSAria to isolate EPCAM^+^CD45^-^DAPI^-^ cells.

#### Single Cell RNA Sequencing and Analysis

B16F10 MTCs extracted from wild type lung, *Prf1^-/-^* lung, and *Prf1^-/-^* bone were hashed using oligonucleotide-conjugated antibodies (TotalSeq B, hashes 1, 2, and 3) to uniquely label individual samples, which were then pooled together in equal proportions to enable multiplexed scRNA-seq. The pooled cells were processed using the 10x Genomics Chromium platform to generate single-cell gel bead-in-emulsions (GEMs). Reverse transcription and cDNA amplification were performed according to the 10x Genomics 3’ v3 protocol. Separate libraries were generated for the gene expression and hashing data. Libraries were sequenced on an Illumina platform to achieve a minimum of 50,000 reads per cell for gene expression and 5,000 reads per cell for hashing. Raw sequencing data were demultiplexed and aligned to the mouse reference genome using Cell Ranger (v6.1.2) from 10x Genomics, which was then used to generate gene expression matrices and hashtag oligonucleotide (HTO) counts for downstream analysis. Cell hashing data were demultiplexed using the Seurat (v4.3.0) HTODemux function to assign sample identity to individual cells. Low-quality cells were filtered based on mitochondrial content (>10%), unique feature counts, and total RNA counts to remove potential doublets and dead cells. Filtered data were normalized and scaled using the SCTransform function in Seurat. Principal component analysis (PCA) was performed, followed by UMAP projection for dimensionality reduction. Clustering was performed using the Louvain algorithm, and clusters were annotated based on canonical marker gene expression. Differential gene expression analysis was conducted between samples using the FindMarkers function in Seurat with the Wilcoxon rank-sum test. Genes with an adjusted p-value < 0.05 were considered statistically significant.

#### Quantitative real-time PCR

Total RNA was isolated from samples using the Qiagen RNeasy Plus Mini Kit according to the manufacturer’s standard protocol. RNA concentrations were quantified with a NanoDrop spectrophotometer. For cDNA synthesis, 2 μg of RNA was reverse transcribed using the Luna One-Step RT-PCR Kit (New England Biolabs) according to the manufacturer’s instructions.

Relative qPCR analysis was performed using GAPDH as the housekeeping gene for normalization and Opn expression as the target gene.

#### Bulk RNA Sequencing

FASTQ files were generated from bulk RNA sequencing and mapped to the UCSC mm10 mouse genome (GTF: Mus_musculus.GRCm38.80) using STAR (v2.5.0a) (*72*). The two-pass alignment method was used, where reads were first aligned using known annotated junctions from Ensembl. Novel junctions identified in the first pass were included in the second pass, during which the RemoveNoncanonical flag was applied. Aligned reads were post-processed with PICARD (v1.124) to add read groups and convert SAM files to sorted, compressed BAM files. Gene expression quantification was performed using HTSeq (v0.5.3) with default parameters. Raw count matrices generated by HTSeq were normalized and analyzed for differential expression using DESeq (R/Bioconductor, v3.2.0). Differential gene expression was assessed using DESeq, applying the Wald test and correcting for multiple testing using the Benjamini-Hochberg method. Genes with an adjusted p-value < 0.05 were considered significant. Normalized log2 expression values were used to perform hierarchical clustering using the Pearson correlation distance metric. Additional dimensionality reduction was performed using multidimensional scaling (MDS) and principal component analysis (PCA). Heatmaps were generated using the heatmap.2 function from the gplots R package, displaying the top 100 differentially expressed genes with mean-centered, normalized log2 expression values. Gene set enrichment analysis (GSEA) was conducted using gene sets from the Broad MSigDB. For groups with fewer than three samples, GSEAPreranked was applied using log2 fold changes generated by DESeq.

#### Patient scRNA-seq samples and data preprocessing

scRNA-seq data were obtained from two cohorts of breast cancer patients treated with immune checkpoint blockade (ICB), including some who received prior chemotherapy. Cells were annotated with metadata including patient ID, treatment status (pre- or post-ICB), and response outcome, as adjudged by the clonal expansion of T cells within the tumor (*52*). Quality control filtering, normalization, and scaling were performed using Seurat (v4.3). Low-quality cells and doublets were excluded based on mitochondrial content, gene counts, and unique feature thresholds. Gene expression data were log-normalized using Seurat’s NormalizeData() function. To assess cytoskeletal regulation, gene lists were compiled from KEGG and GSEA sources (Table 1). Module scores were calculated using Seurat’s AddModuleScore() function across the full dataset as well as a subset of pre-treatment cells. These scores were used to evaluate pathway-level expression in relation to ICB treatment outcome. Module score distributions were visualized using violin plots. To evaluate gene expression of actin cytoskeletal genes compared to intermediate filament and microtubule genes, curated gene lists were generated using literature sources (Table 2). Dot plots and heatmaps were used to summarize averaged gene expression per outcome group. Feature plots were generated with consistent scaling across panels for visual comparability. Statistical significance between groups was assessed using Wilcoxon rank-sum tests unless otherwise stated.

#### Statistics

Analyses were carried out using either representative experiments or pooled data as indicated (*n* is defined in the figure legends for each experiment). Statistical tests (two-tailed Mann-Whitney, two-tailed ANOVA, paired and unpaired two-tailed t-tests and Log-rank Mantel-Cox tests) were performed using GraphPad Prism. Unless otherwise indicated, error bars denote SEM. No statistical methods were used to determine sample size prior to experiments.

---

**Fig. S1:**
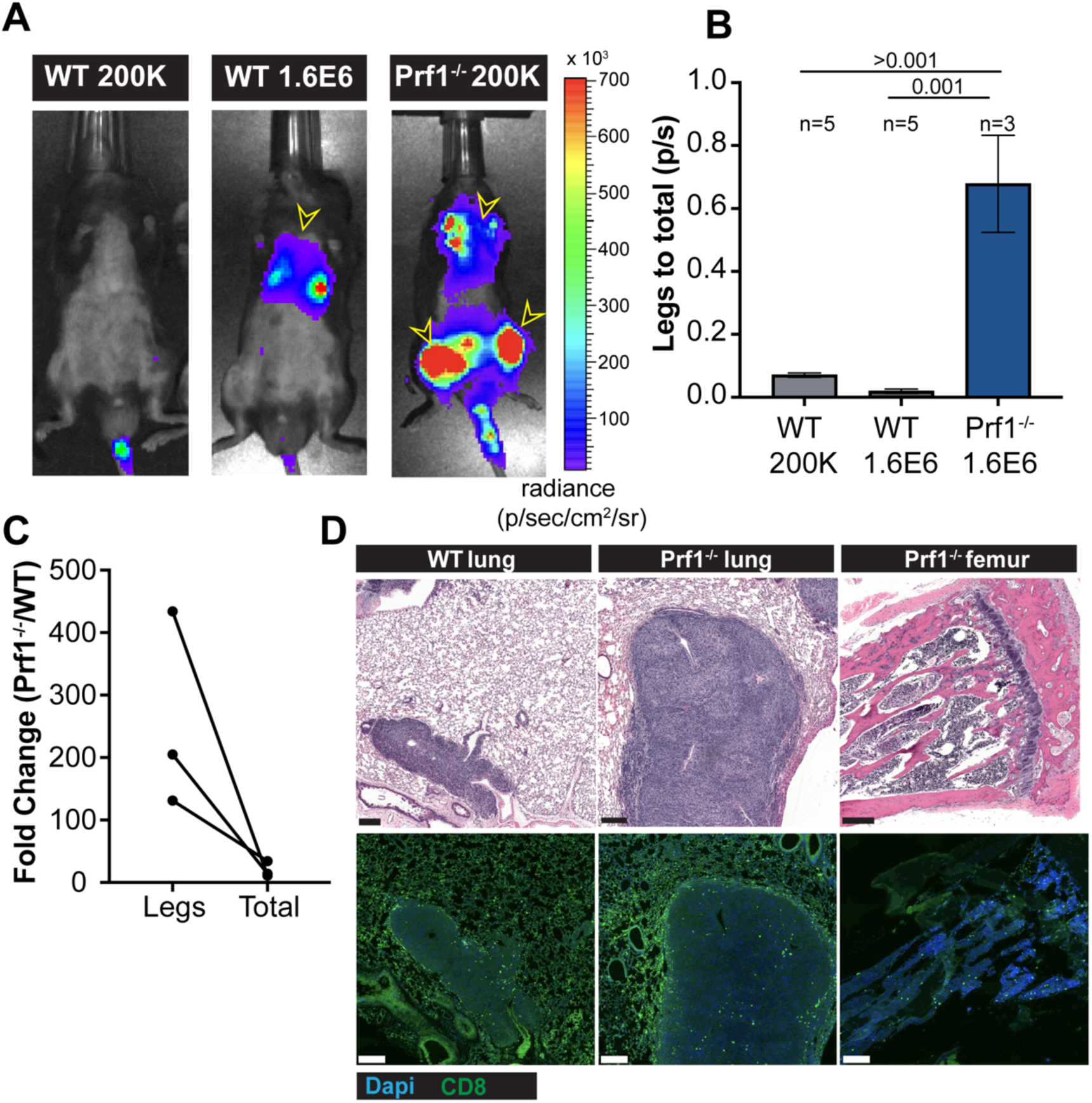
Preferential suppression of bone metastasis by cytotoxic lymphocytes. (A-B) Wild type and *Prf1^-/-^* mice were i.v. injected with low (2 × 10^5^) or high (1.6 × 10^6^) doses of Luc^+^ B16F10 cells as indicated, and the mice were monitored for metastatic colonization of the lungs and bone. (A) Representative IVIS images of tumor-bearing wild type and *Prf1^-/-^* mice, with metastatic burden in the lungs and femurs indicated by black and yellow arrowheads. (B) Quantification of relative femoral colonization, expressed as a ratio of IVIS signal in the legs to the total IVIS signal. Error bars denote SEM. Sample size is indicated above each bar. P-values calculated by one-way ANOVA. (C) Mean fold change in B16F10 colonization in *Prf1^-/-^* mice relative to wild type controls, determined for total metastasis and femoral metastasis (legs). Paired values were derived from 3 independent experiments. (D) H&E images of representative metastatic tumors from the indicated tissues are shown above, with immunofluorescence staining of CD8^+^ T cells shown below. Scale bars = 200 μm.

**Fig. S2:**
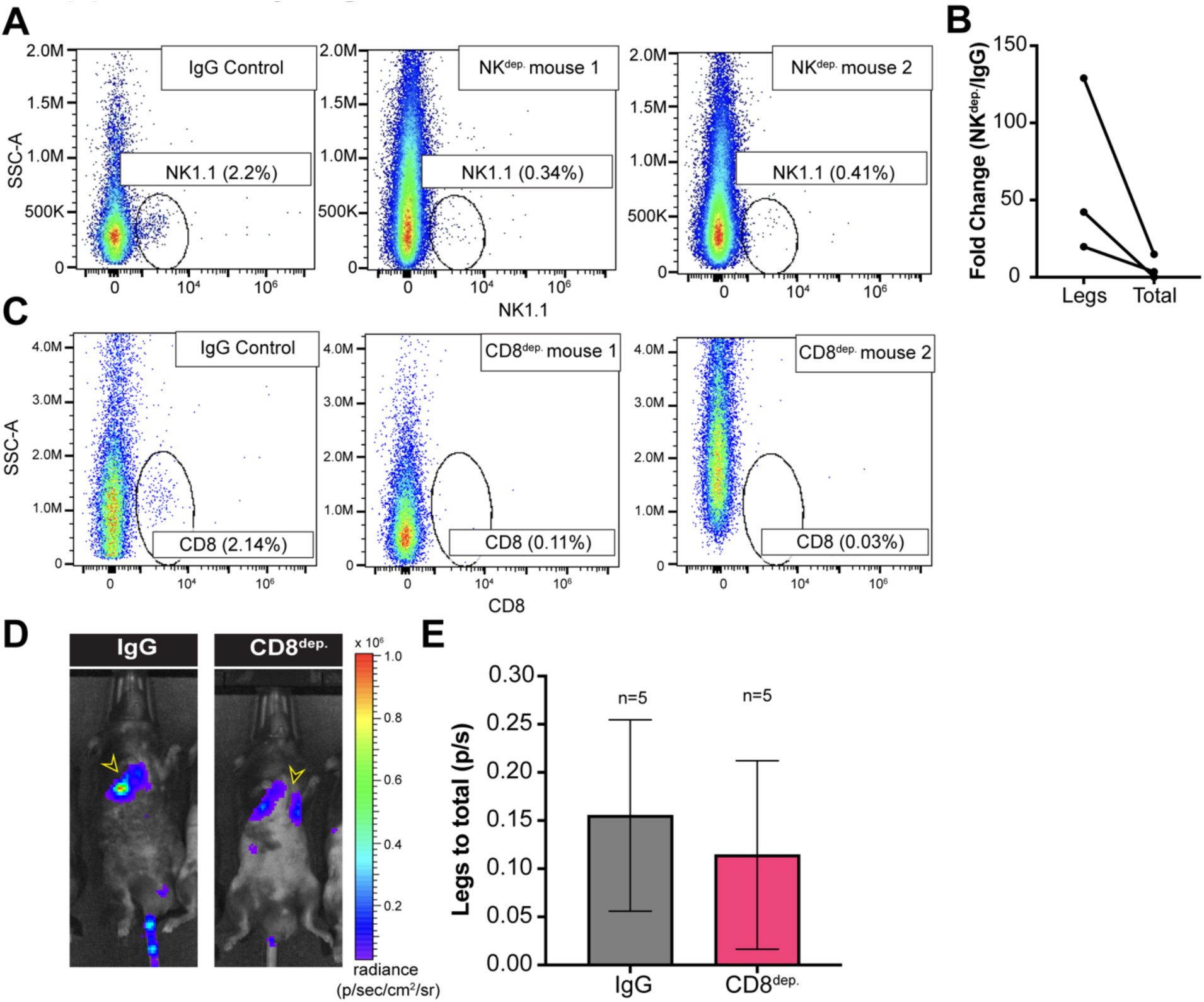
Depletion of NK cells, but not CD8^+^ T cells, alters B16F10 metastatic site distribution. (A) Flow cytometric validation of NK cell depletion by anti-asialoGM1 antibody treatment. NK1.1 staining of blood samples from representative IgG control and NK depleted (NK^dep.^) mice are shown. (B) Mean fold change in B16F10 colonization in NK^dep.^ mice relative to IgG controls, determined for total metastasis and femoral metastasis (legs). Paired values were derived from 3 independent experiments. (C) Flow cytometric validation of CD8^+^ T cell depletion by anti-CD8 antibody treatment. CD8 staining of blood samples from representative IgG control and CD8 depleted (CD8^dep.^) mice are shown. (D-E) Luc^+^ B16F10 cells were injected i.v. into IgG control and CD8^dep.^ recipient mice, which were then monitored for metastatic colonization of the lungs and bone. (D) Representative IVIS images of tumor-bearing IgG control and CD8^dep.^ mice, with metastatic burden in the lungs and femurs indicated by black and yellow arrowheads. (E) Quantification of relative femoral colonization, expressed as a ratio of IVIS signal in the legs to the total IVIS signal. Error bars denote SEM. Sample size is indicated above each bar. P-value calculated by unpaired Student’s t-test.

**Fig. S3:**
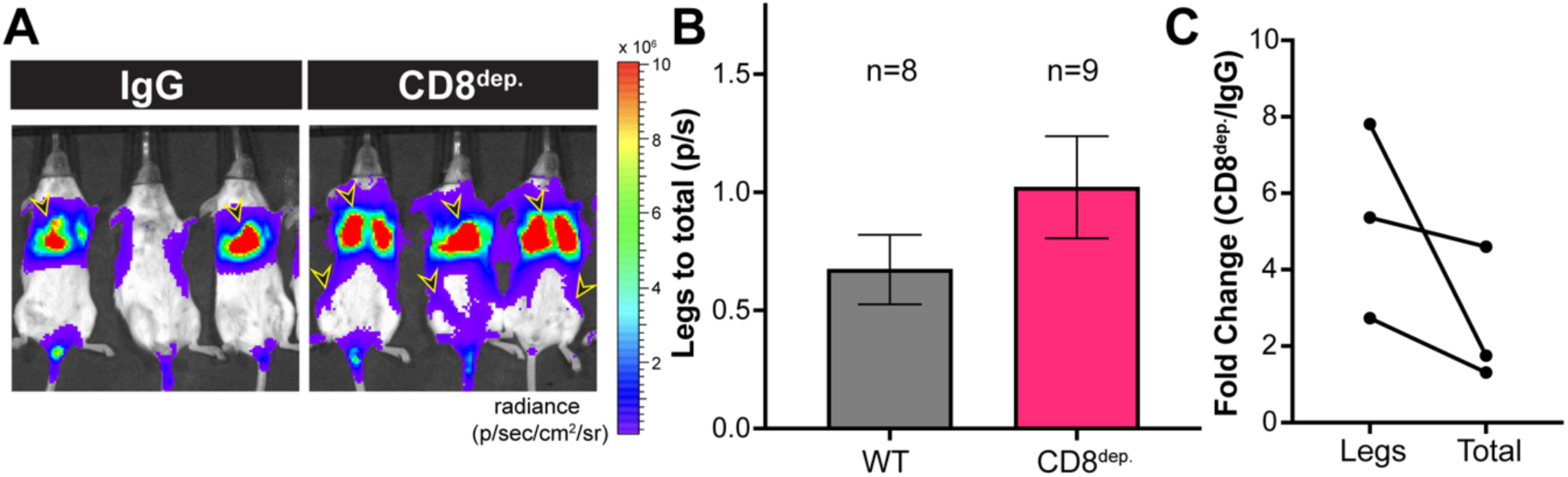
Preferential suppression of EMT6 bone metastasis by CD8^+^ T cells. Luc^+^ EMT6 cells were injected i.v. into IgG control and CD8^dep.^ recipient mice, which were then monitored for metastatic colonization of the lungs and bone. (A) Representative IVIS images of tumor-bearing IgG control and CD8^dep.^ mice, with metastatic burden in the lungs and femurs indicated by black and yellow arrowheads. (B) Quantification of relative femoral colonization, expressed as a ratio of IVIS signal in the legs to the total IVIS signal. Error bars denote standard error of the mean (SEM). Sample size is indicated above each bar. (C) Mean fold change in EMT6 colonization in CD8^dep.^ mice relative to wild type controls, determined for total metastasis and femoral metastasis (legs). Paired values were derived from 3 independent experiments.

**Fig. S4:**
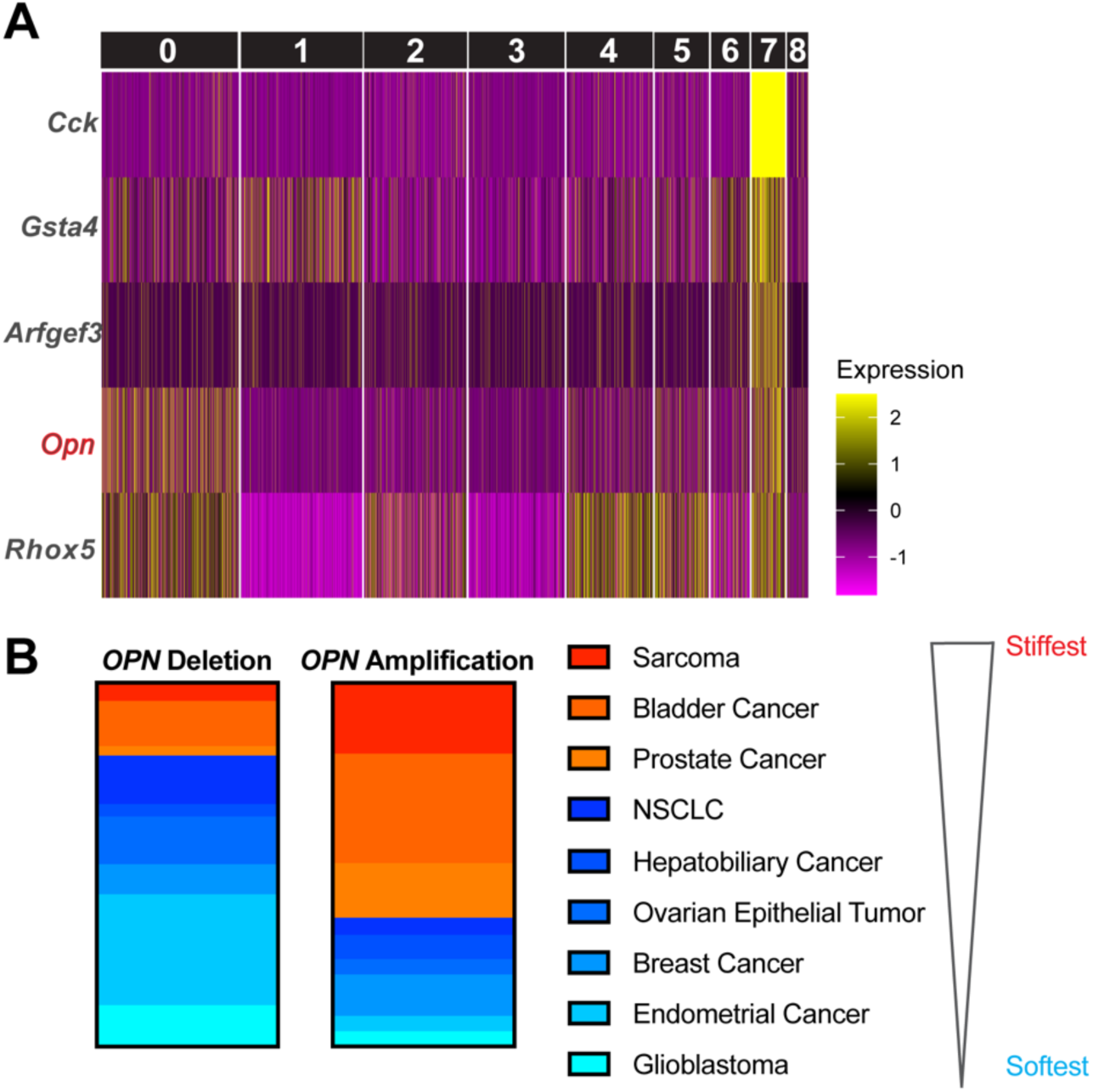
Transcriptional features of bone metastases. (A) GFP^+^ B16F10 cells were i.v. injected either into wild type or *Prf1^-/-^* mice, and after 2 weeks, MTCs from the resulting metastases in the lungs of wild type mice and the lungs and bones of *Prf1^-/-^* mice were extracted and subjected to scRNA-seq. Heatmap shows the top 5 differentially expressed genes in cluster 7 (see Fig. 4). (B) Graph showing the relative frequency of *OPN* amplification and deletion in human tumors, determined using TCGA data. Tumors are organized according to the rigidity of their parent tissue.

**Fig. S5:**
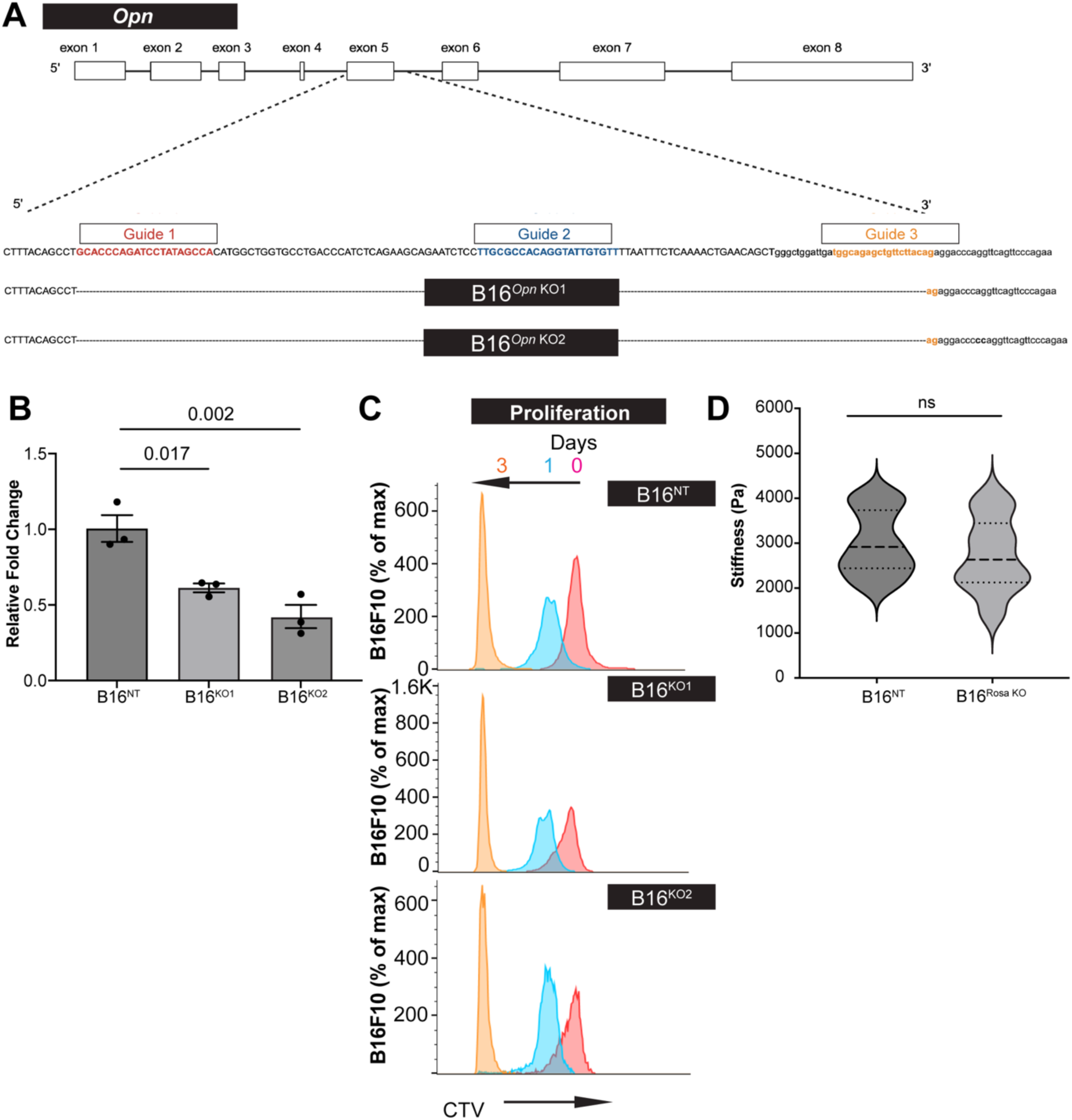
CRISPR/Cas9 targeting of *Opn* in B16F10 cells. (A) Schematic diagram of the CRISPR/Cas9 multi-guide approach used to knock out *Opn* in B16F10 cells. The exon configuration of the gene is shown above and the targeted locus below, along with the 134 bp and 132 bp deletions induced in Opn-KO1 and Opn-KO2, respectively. (B) Opn expression in the indicated NT and Opn-KO cell lines, determined by qRT-PCR. Error bars denote SEM of 3 technical replicates. P-values determined by one-way ANOVA. (C) The indicated B16F10 cell lines were stained with cell trace violet (CTV), and their proliferation assessed by dye dilution. Representative flow cytometric measurements of CTV fluorescence at 0, 1, and 3 days of culture are shown. (D) AFM stiffness measurements of NT B16F10 cells and Rosa knockout “safe harbor” B16F10 controls. violins encompass the entire data distribution, with dashed lines denoting the median and dotted lines indicating the upper and lower quartiles. P-value calculated by unpaired Student’s t-test.

**Fig. S6:**
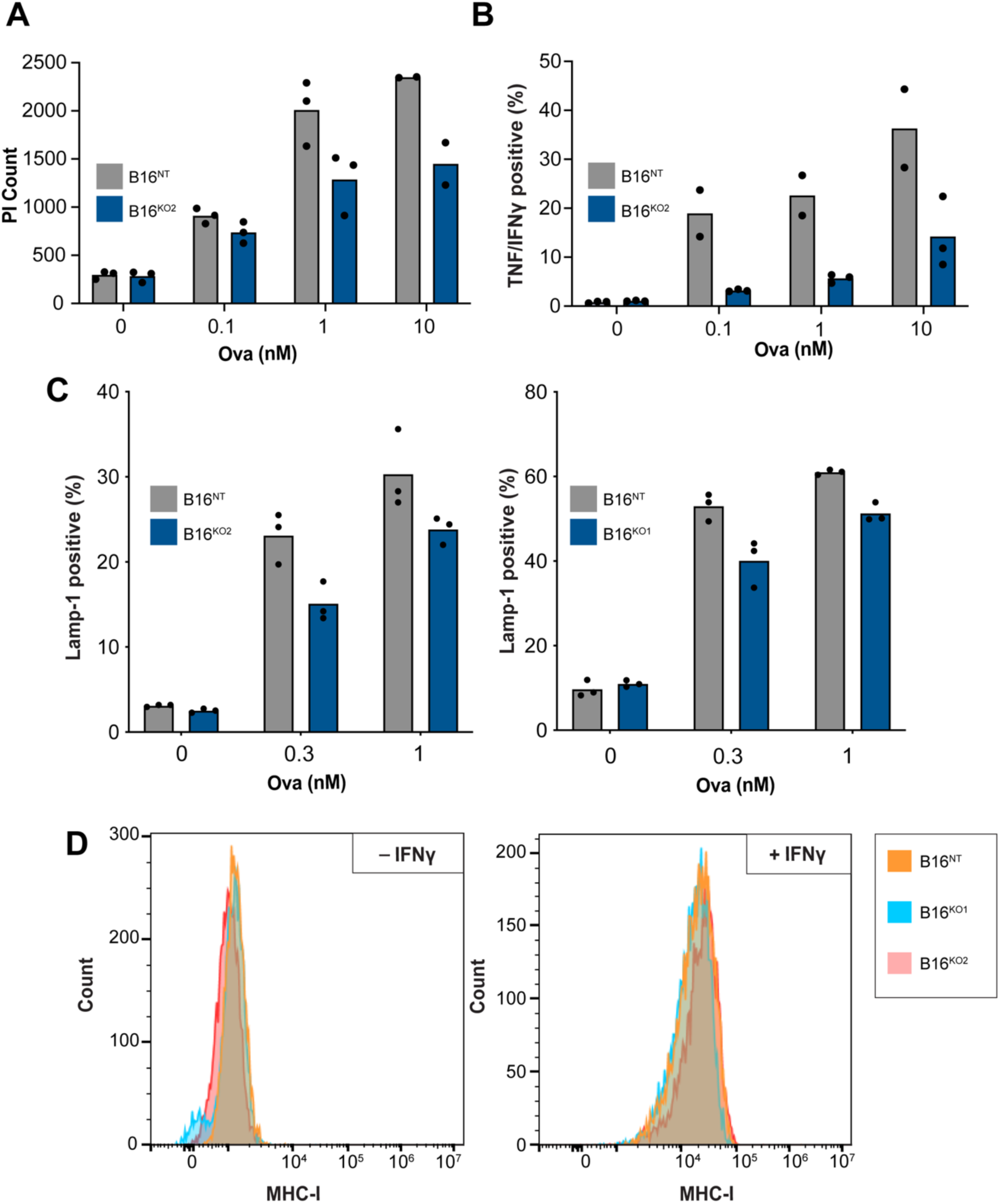
Opn sensitizes B16F10 cells to cytotoxic lymphocytes. The indicated B16F10 cells were loaded with increasing amounts of OVA and then mixed with OT-1 CTLs. (A) Target cell killing, measured by PI influx after 5 h. (B) CTL cytokine production, measured by intracellular staining for TNF and IFNγ, after 5 h. (C) Degranulation, measured by surface exposure of Lamp1, after 90 min. Opn-KO2 results are shown on the left, and Opn-KO1 results on the right. (D) Representative histograms showing MHC-I expression on the indicated B16F10 cells lines, determined in the absence (left) or presence (right) of 20 ng/mL IFNγ, which promotes upregulation of class I MHC (MHC-I).

**Fig. S7:**
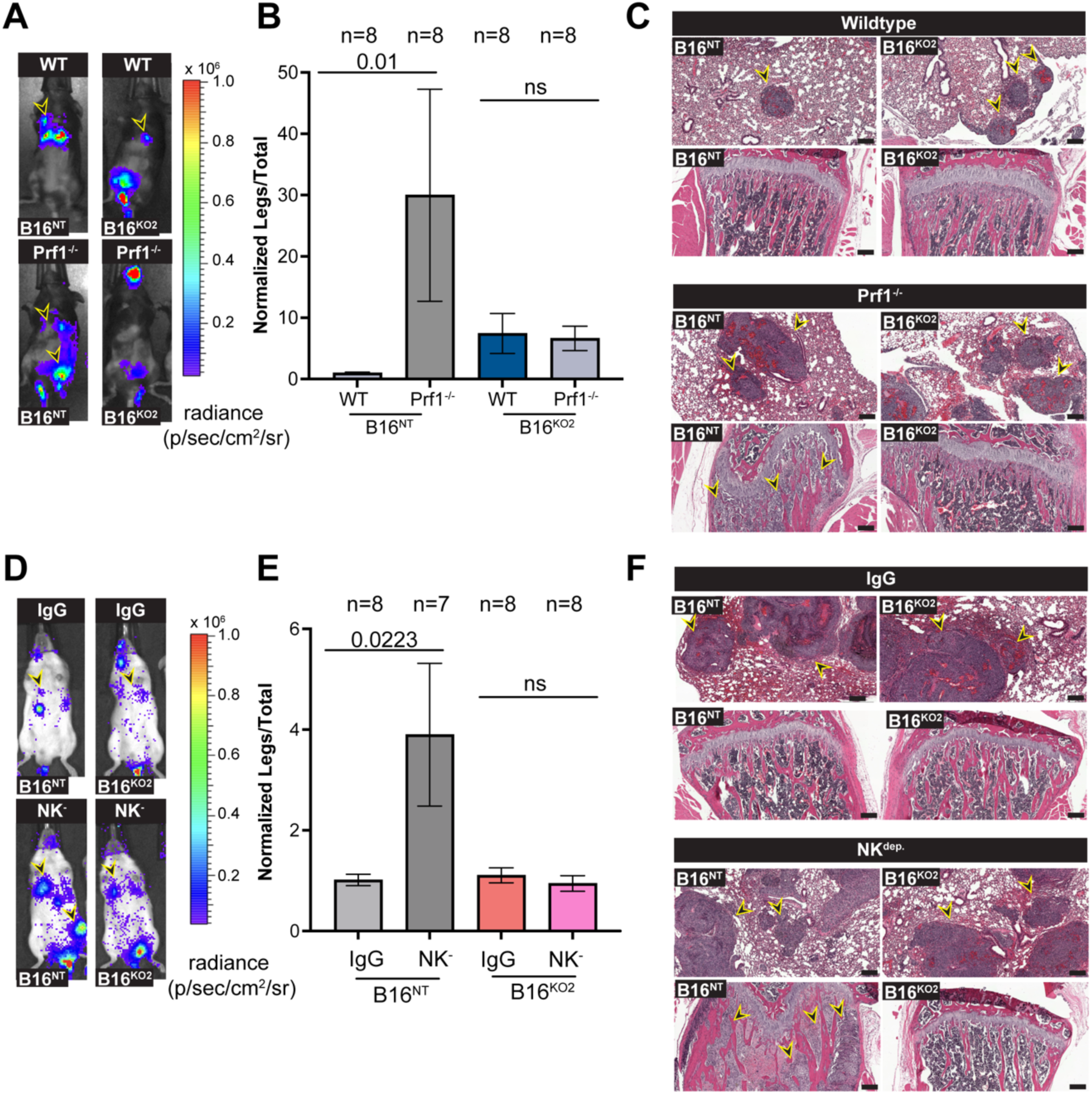
Opn is required for mechanoreciprocity *in vivo*. (A-C) Luc^+^ NT or Opn-KO2 B16F10 cells were injected i.v. into wild type and *Prf1^-/-^* mice, which were then monitored for metastatic colonization of the lungs and bone. (A) Representative IVIS images of tumor-bearing wild type and *Prf1^-/-^* mice injected with the indicated B16F10 lines, with metastatic burden in the lungs and femurs indicated by black and yellow arrowheads. (B) Quantification of relative femoral colonization, expressed as a ratio of IVIS signal in the legs to the total IVIS signal. (C) Representative H&E sections highlighting the absence of femoral colonization in *Prf1^-/-^*mice injected with Opn-KO2 B16F10 cells. Lung metastases and B16-NT metastases in *Prf1^-/-^* bone are indicated by black and yellow arrowheads. Scale bars = 200 μm. (D-F) Luc^+^ NT or Opn-KO2 B16F10 cells were injected i.v. into IgG control and NK^dep.^ recipient mice, which were then monitored for metastatic colonization of the lungs and bone. (D) Representative IVIS images of tumor-bearing IgG control and NK dep mice injected with the indicated B16F10 lines, with metastatic burden in the lungs and femurs indicated by black and yellow arrowheads. (E) Quantification of relative femoral colonization, expressed as a ratio of IVIS signal in the legs to the total IVIS signal. (F) Representative H&E sections highlighting the absence of femoral colonization in NK^dep.^ mice injected with Opn-KO2 B16F10 cells. Lung metastases and B16-NT metastases in NK^dep.^ bone are indicated by black and yellow arrowheads. Scale bars = 200 μm. In B and E, error bars denote SEM, sample size is indicated above each bar, and P-values were calculated by one-way ANOVA.

**Fig. S8:**
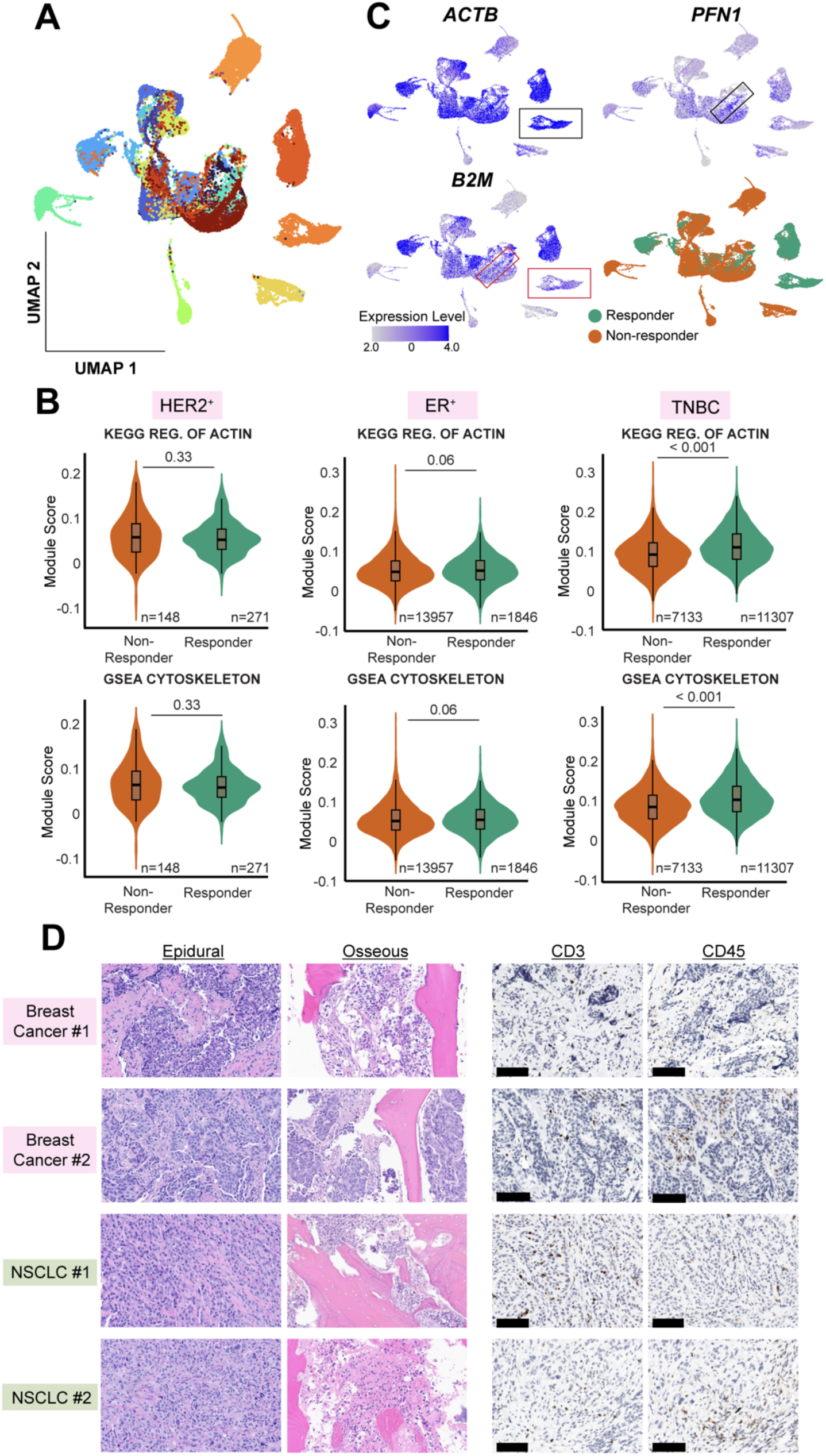
Transcriptomic and histological analyses of patient tumors. (A-B) scRNA-seq analysis of cytoskeletal gene expression in breast cancer patients treated with ICB. Breast cancer biopsies were collected before and 9-11 days after a single dose of anti-PD-1 immunotherapy, then subjected to scRNA-seq. (A) UMAP visualization of scRNA-seq data using only pre-treatment samples, colored by patient ID. (B) F-actin cytoskeleton gene expression in non-responder versus responder tumors, calculated using data from pre-treatment samples of the indicated breast cancer classes. Module scores were generated using the KEGG “Regulation of Actin Cytoskeleton” pathway and a GSEA “Cytoskeleton” gene set. Embedded boxes indicate median and interquartile range. P-values calculated by Wilcoxon rank-sum test. (C) UMAP feature plots showing expression of *ACTB*, *PFN1*, and *B2M*. The same UMAP plot colored by response status (responder versus non-responder) is shown for reference. Boxed regions highlight responder cells with relatively low *B2M* expression but high expression of *ACTB* or *PFN1*. (D) Histological analysis of spinal metastases. Left, H&E staining of representative sections from epidural and osseous regions of the indicated tumors. Right, IHC staining of CD3 and CD45 in representative epidural sections of the indicated tumors. Scale bars = 100 μm.

**Fig. S9:**
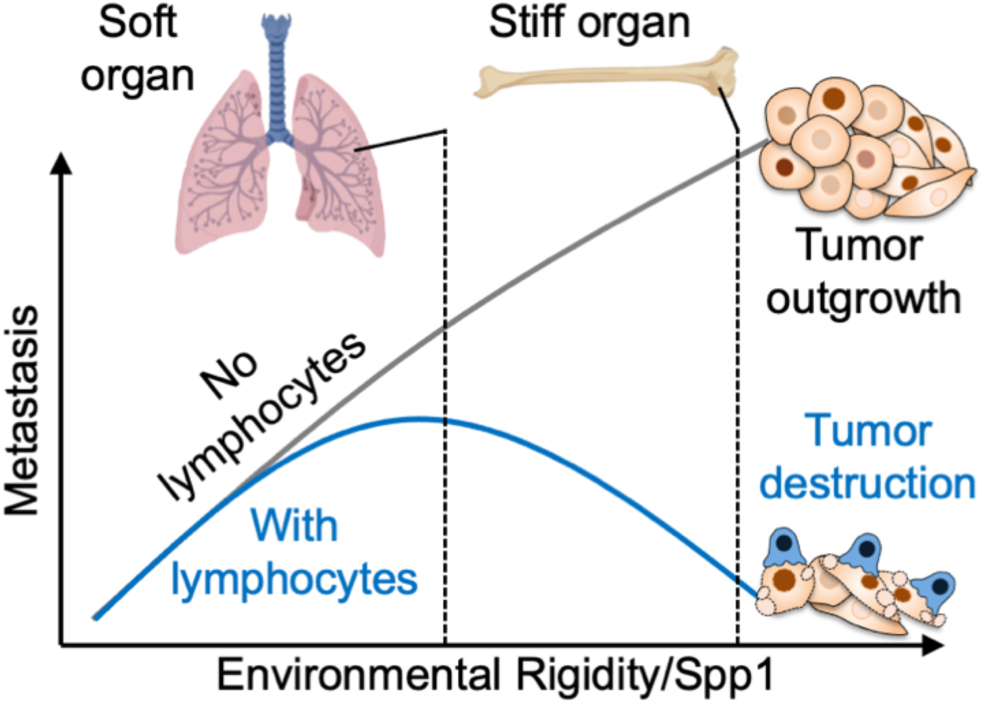
Model demonstrating how the combined effects of mechanoreciprocity and mechanosurveillance determine metastatic site distribution. Mechanoreciprocity enables MTCs to colonize rigid environments like the bone, but it also stiffens the MTCs, which sensitizes them to destruction by cytotoxic lymphocytes. This manifests in the preferential suppression of bone metastasis in immunocompetent settings.

**Table S1:**
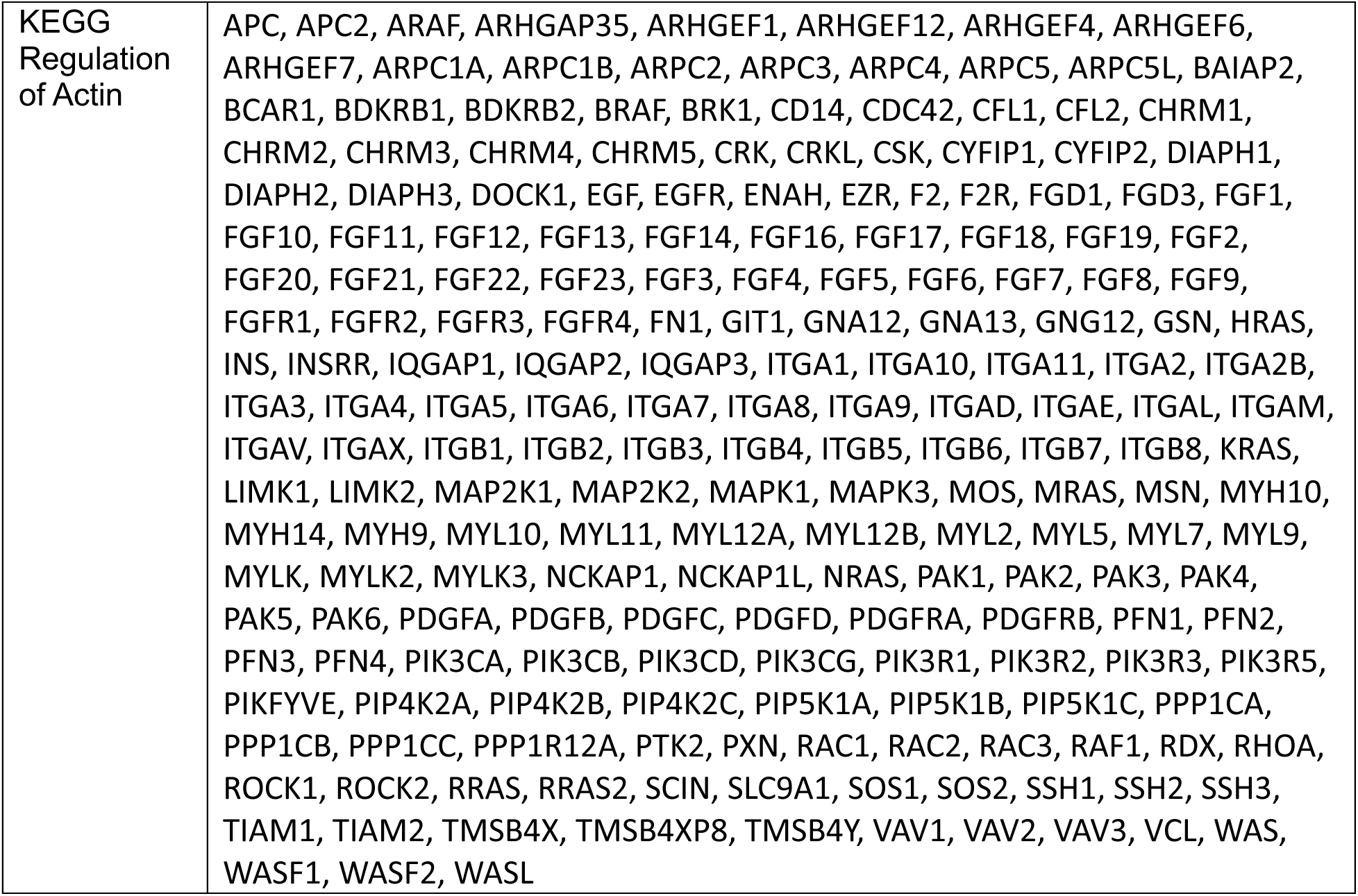
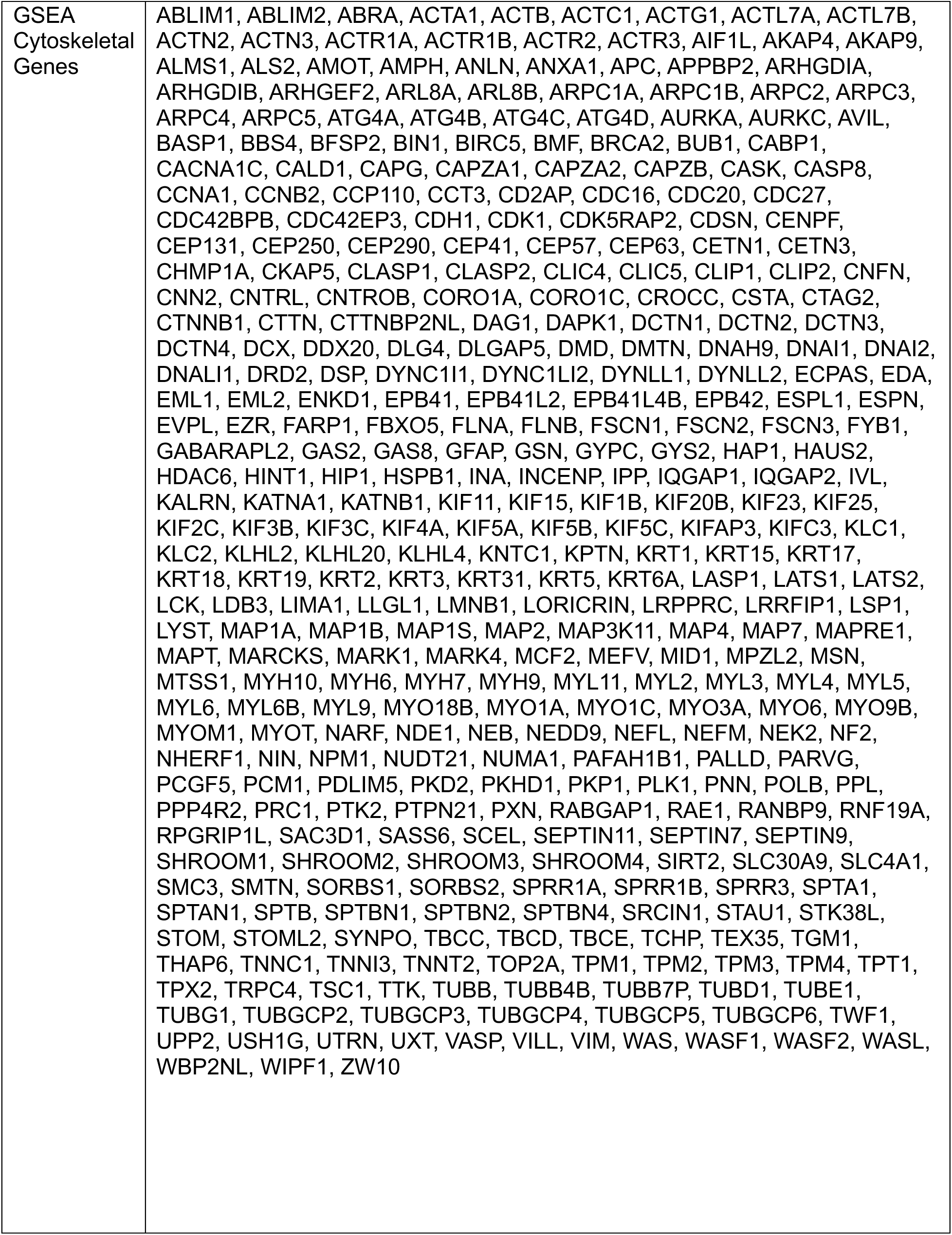
Cytoskeletal gene lists.

**Table S2:**
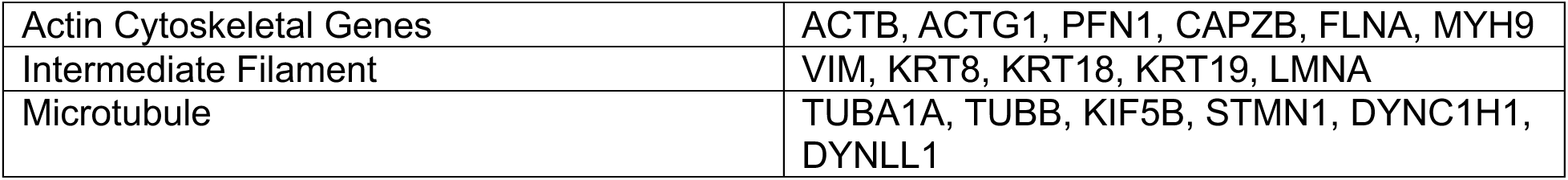
Curated gene lists for actin cytoskeletal genes, intermediate filament related genes, and microtubule related genes.

## References and Notes

1. G. P. Gupta, J. Massague, Cancer metastasis: building a framework. Cell 127, 679–695 (2006).

2. D. X. Nguyen, P. D. Bos, J. Massague, Metastasis: from dissemination to organ-specific colonization. Nat Rev Cancer 9, 274–284 (2009).

3. J. Fares, M. Y. Fares, H. H. Khachfe, H. A. Salhab, Y. Fares, Molecular principles of metastasis: a hallmark of cancer revisited. Signal Transduct Target Ther 5, 28 (2020).

4. T. N. Seyfried, L. C. Huysentruyt, On the origin of cancer metastasis. Crit Rev Oncog 18, 43–73 (2013).

5. C. F. Guimaraes, L. Gasperini, A. P. Marques, R. L. Reis, The stiffness of living tissues and its implications for tissue engineering. Nat Rev Mater 5, 351–370 (2020).

6. T. Iskratsch, H. Wolfenson, M. P. Sheetz, Appreciating force and shape-the rise of mechanotransduction in cell biology. Nat Rev Mol Cell Biol 15, 825–833 (2014).

7. M. J. Paszek, V. M. Weaver, The tension mounts: mechanics meets morphogenesis and malignancy. J Mammary Gland Biol Neoplasia 9, 325–342 (2004).

8. D. T. Butcher, T. Alliston, V. M. Weaver, A tense situation: forcing tumour progression. Nat Rev Cancer 9, 108–122 (2009).

9. A. Agrawal et al., Mechanical signatures in cancer metastasis. NPJ Biol Phys Mech 2, 3 (2025).

10. J. Solon, I. Levental, K. Sengupta, P. C. Georges, P. A. Janmey, Fibroblast adaptation and stiffness matching to soft elastic substrates. Biophysical Journal 93, 4453–4461 (2007).

11. F. J. Byfield, R. K. Reen, T. P. Shentu, I. Levitan, K. J. Gooch, Endothelial actin and cell stiffness is modulated by substrate stiffness in 2D and 3D. J Biomech 42, 1114–1119 (2009).

12. M. Guo et al., Cell volume change through water efflux impacts cell stiffness and stem cell fate. P Natl Acad Sci USA 114, E8618–E8627 (2017).

13. R. J. Pelham, Jr., Y. Wang, Cell locomotion and focal adhesions are regulated by substrate flexibility. Proc Natl Acad Sci U S A 94, 13661–13665 (1997).

14. O. J. Finn, A Believer’s Overview of Cancer Immunosurveillance and Immunotherapy. J Immunol 200, 385–391 (2018).

15. L. M. E. Janssen, E. E. Ramsay, C. D. Logsdon, W. W. Overwijk, The immune system in cancer metastasis: friend or foe? J Immunother Cancer 5, 79 (2017).

16. N. Xie et al., Neoantigens: promising targets for cancer therapy. Signal Transduct Target Ther 8, 9 (2023).

17. L. L. Lanier, NK cell recognition. Annu Rev Immunol 23, 225–274 (2005).

18. M. L. Dustin, E. O. Long, Cytotoxic immunological synapses. Immunol Rev 235, 24–34 (2010).

19. S. C. Wei, C. R. Duffy, J. P. Allison, Fundamental Mechanisms of Immune Checkpoint Blockade Therapy. Cancer Discov 8, 1069–1086 (2018).

20. R. Basu, M. Huse, Mechanical Communication at the Immunological Synapse. Trends Cell Biol 27, 241–254 (2017).

21. W. Chen, J. Lou, C. Zhu, Forcing switch from short- to intermediate- and long-lived states of the alphaA domain generates LFA-1/ICAM-1 catch bonds. J Biol Chem 285, 35967–35978 (2010).

22. C. Gonzalez et al., Nanobody-CD16 Catch Bond Reveals NK Cell Mechanosensitivity. Biophys J 116, 1516–1526 (2019).

23. B. Liu, W. Chen, B. D. Evavold, C. Zhu, Accumulation of dynamic catch bonds between TCR and agonist peptide-MHC triggers T cell signaling. Cell 157, 357–368 (2014).

24. D. Blumenthal, V. Chandra, L. Avery, J. K. Burkhardt, Mouse T cell priming is enhanced by maturation-dependent stiffening of the dendritic cell cortex. Elife 9, e55995 (2020).

25. E. Judokusumo, E. Tabdanov, S. Kumari, M. L. Dustin, L. C. Kam, Mechanosensing in T lymphocyte activation. Biophys J 102, L5–7 (2012).

26. M. Saitakis et al., Different TCR-induced T lymphocyte responses are potentiated by stiffness with variable sensitivity. Elife 6, e23190 (2017).

27. Z. Wan et al., B cell activation is regulated by the stiffness properties of the substrate presenting the antigens. J Immunol 190, 4661–4675 (2013).

28. M. Tello-Lafoz et al., Cytotoxic lymphocytes target characteristic biophysical vulnerabilities in cancer. Immunity 54, 1037–1054 e1037 (2021).

29. K. Lei et al., Cancer-cell stiffening via cholesterol depletion enhances adoptive T-cell immunotherapy. Nat Biomed Eng 5, 1411–1425 (2021).

30. Y. Liu et al., Cell Softness Prevents Cytolytic T-cell Killing of Tumor-Repopulating Cells. Cancer Res 81, 476–488 (2021).

31. J. H. Kang et al., Noninvasive monitoring of single-cell mechanics by acoustic scattering (vol 16, pg 263, 2019). Nature Methods 16, 270-270 (2019).

32. C. Khanna, K. Hunter, Modeling metastasis in vivo. Carcinogenesis 26, 513–523 (2005).

33. W. E. Seaman, M. Sleisenger, E. Eriksson, G. C. Koo, Depletion of natural killer cells in mice by monoclonal antibody to NK-1.1. Reduction in host defense against malignancy without loss of cellular or humoral immunity. J Immunol 138, 4539–4544 (1987).

34. J. Hsu et al., Contribution of NK cells to immunotherapy mediated by PD-1/PD-L1 blockade. Journal of Clinical Investigation 128, 4654–4668 (2018).

35. R. Piranlioglu et al., Primary tumor-induced immunity eradicates disseminated tumor cells in syngeneic mouse model. Nat Commun 10, 1430 (2019).

36. R. A. Carpenter, J. G. Kwak, S. R. Peyton, J. Lee, Implantable pre-metastatic niches for the study of the microenvironmental regulation of disseminated human tumour cells. Nat Biomed Eng 2, 915–929 (2018).

37. A. Bastos et al., The Intracellular and Secreted Sides of Osteopontin and Their Putative Physiopathological Roles. Int J Mol Sci 24, (2023).

38. Y. Asou et al., Osteopontin facilitates angiogenesis, accumulation of osteoclasts, and resorption in ectopic bone. Endocrinology 142, 1325–1332 (2001).

39. M. A. Chellaiah et al., Osteopontin deficiency produces osteoclast dysfunction due to reduced CD44 surface expression. Mol Biol Cell 14, 173–189 (2003).

40. A. W. Watson et al., Breast tumor stiffness instructs bone metastasis via maintenance of mechanical conditioning. Cell Rep 35, 109293 (2021).

41. Y. You et al., Higher Matrix Stiffness Upregulates Osteopontin Expression in Hepatocellular Carcinoma Cells Mediated by Integrin beta1/GSK3beta/beta-Catenin Signaling Pathway. PLoS One 10, e0134243 (2015).

42. Y. Kang et al., A multigenic program mediating breast cancer metastasis to bone. Cancer Cell 3, 537–549 (2003).

43. S. D. Bhattacharya et al., Osteopontin regulates epithelial mesenchymal transition-associated growth of hepatocellular cancer in a mouse xenograft model. Ann Surg 255, 319–325 (2012).

44. M. Kovacheva, M. Zepp, M. Schraad, S. Berger, M. R. Berger, Conditional Knockdown of Osteopontin Inhibits Breast Cancer Skeletal Metastasis. Int J Mol Sci 20, (2019).

45. Y. Zhao et al., Tumor alphavbeta3 integrin is a therapeutic target for breast cancer bone metastases. Cancer Res 67, 5821–5830 (2007).

46. H. Nemoto et al., Osteopontin deficiency reduces experimental tumor cell metastasis to bone and soft tissues. J Bone Miner Res 16, 652–659 (2001).

47. S. J. Hotte, E. W. Winquist, L. Stitt, S. M. Wilson, A. F. Chambers, Plasma osteopontin: associations with survival and metastasis to bone in men with hormone-refractory prostate carcinoma. Cancer 95, 506–512 (2002).

48. X. Pang et al., SPP1 Promotes Enzalutamide Resistance and Epithelial-Mesenchymal-Transition Activation in Castration-Resistant Prostate Cancer via PI3K/AKT and ERK1/2 Pathways. Oxid Med Cell Longev 2021, 5806602 (2021).

49. X. Hou et al., Osteopontin is a useful predictor of bone metastasis and survival in patients with locally advanced nasopharyngeal carcinoma. Int J Cancer 137, 1672–1678 (2015).

50. H. Singhal et al., Elevated plasma osteopontin in metastatic breast cancer associated with increased tumor burden and decreased survival. Clin Cancer Res 3, 605–611 (1997).

51. G. Carlinfante et al., Differential expression of osteopontin and bone sialoprotein in bone metastasis of breast and prostate carcinoma. Clin Exp Metastasis 20, 437–444 (2003).

52. A. Bassez et al., A single-cell map of intratumoral changes during anti-PD1 treatment of patients with breast cancer. Nat Med 27, 820–832 (2021).

53. I. A. Vathiotis et al., Immune Checkpoint Blockade in Hormone Receptor-Positive Breast Cancer: Resistance Mechanisms and Future Perspectives. Clin Breast Cancer 22, 642–649 (2022).

54. L. Y. Dirix et al., Avelumab, an anti-PD-L1 antibody, in patients with locally advanced or metastatic breast cancer: a phase 1b JAVELIN Solid Tumor study. Breast Cancer Res Treat 167, 671–686 (2018).

55. K. Dhatchinamoorthy, J. D. Colbert, K. L. Rock, Cancer Immune Evasion Through Loss of MHC Class I Antigen Presentation. Front Immunol 12, 636568 (2021).

56. L. B. Alexandrov et al., Signatures of mutational processes in human cancer. Nature 500, 415–421 (2013).

57. J. Zhang, D. Huang, P. E. Saw, E. Song, Turning cold tumors hot: from molecular mechanisms to clinical applications. Trends Immunol 43, 523–545 (2022).

58. S. Gan et al., Distinct tumor architectures and microenvironments for the initiation of breast cancer metastasis in the brain. Cancer Cell 42, 1693–1712 e1624 (2024).

59. J. Ning et al., Macrophage-coated tumor cluster aggravates hepatoma invasion and immunotherapy resistance via generating local immune deprivation. Cell Rep Med 5, 101505 (2024).

60. B. Qian et al., A distinct macrophage population mediates metastatic breast cancer cell extravasation, establishment and growth. PLoS One 4, e6562 (2009).

61. F. S. Majedi et al., T-cell activation is modulated by the 3D mechanical microenvironment. Biomaterials 252, 120058 (2020).

62. E. E. Er, M. Tello-Lafoz, M. Huse, Mechanoregulation of Metastasis beyond the Matrix. Cancer Res 82, 3409–3419 (2022).

63. S. Nasrollahi et al., Past matrix stiffness primes epithelial cells and regulates their future collective migration through a mechanical memory. Biomaterials 146, 146–155 (2017).

64. C. Yang, M. W. Tibbitt, L. Basta, K. S. Anseth, Mechanical memory and dosing influence stem cell fate. Nat Mater 13, 645–652 (2014).

65. S. Choi et al., Bone-matrix mineralization dampens integrin-mediated mechanosignalling and metastatic progression in breast cancer. Nat Biomed Eng 7, 1455–1472 (2023).

66. F. Kai, A. P. Drain, V. M. Weaver, The Extracellular Matrix Modulates the Metastatic Journey. Dev Cell 49, 332–346 (2019).

67. T. Ibrahim et al., Expression of bone sialoprotein and osteopontin in breast cancer bone metastases. Clin Exp Metastasis 18, 253–260 (2000).

68. J. D. Klement et al., An osteopontin/CD44 immune checkpoint controls CD8+ T cell activation and tumor immune evasion. J Clin Invest 128, 5549–5560 (2018).

69. J. D. Klement et al., Osteopontin Blockade Immunotherapy Increases Cytotoxic T Lymphocyte Lytic Activity and Suppresses Colon Tumor Progression. Cancers (Basel) 13, (2021).

70. E. E. Er et al., Pericyte-like spreading by disseminated cancer cells activates YAP and MRTF for metastatic colonization. Nat Cell Biol 20, 966–978 (2018).

71. J. Lee et al., Implantable microenvironments to attract hematopoietic stem/cancer cells. Proc Natl Acad Sci U S A 109, 19638–19643 (2012).

72. A. Dobin et al., STAR: ultrafast universal RNA-seq aligner. Bioinformatics 29, 15–21 (2013).

